# Falling shoulders ahead: adaptive alien genes are maintained amid vanishing introgression footprint in sea squirts

**DOI:** 10.1101/2023.09.21.558831

**Authors:** Fanny Touchard, Frédérique Cerqueira, Nicolas Bierne, Frédérique Viard

## Abstract

Human transport of species across oceans disrupts natural dispersal barriers and facilitates hybridisation between previously allopatric species. The recent introduction of the North Pacific sea squirt, *Ciona robusta*, into the native range of the North Atlantic sea squirt, *C. intestinalis*, is a good example of this outcome. Recent studies have revealed an adaptive introgression, in a single chromosomal region, from the introduced into the native species. Here, we monitored this adaptive introgression over time, examining both the frequency of adaptive alien genes at the core and the hitchhiking footprint in the shoulders of the introgression island, by studying a thousand *Ciona spp.* individuals collected in 22 ports of the contact zone, 14 of which were sampled 20 generations apart. For that purpose, we developed a KASP multiplex genotyping approach, which proved effective in identifying native, non-indigenous and hybrid individuals and in detecting introgressed haplotypes. Adaptive alien genes were detected where they had been found 20 generations ago, as well as in some newly sampled ports. The core region of the introgression, where the frequency of *C. robusta* alleles is the highest and locally adaptive genes must be, remains stable in both space and time. In contrast, we observed an erosion of *C. robusta* ancestry tracts in flanking chromosomal shoulders on the edges of the core, consistent with the second phase of a local sweep and a purge of incompatible introgressed alleles. Our study reveals how the shoulders of an adaptive introgression island fall over time after the initial sweep.

## 1. Introduction

Human activities are responsible for ever-increasing biological introductions (Seebens et al. 2017), and the marine environment is no exception (Costello et al., 2022; de Castro et al., 2017; Hulme, 2021; Sardain et al., 2019; Zenetos et al. 2022). Anthropogenic transport of species facilitates secondary contact between previously isolated species or lineages by altering connectivity pathways and acting as corridors and stepping stones (Airoldi et al., 2015; Alter et al., 2020; Bishop et al., 2017). These secondary contacts can result in anthropogenic hybridisations between native and non-indigenous species or lineages, and, as are post-glacial hybrid zones (Hewitt, 1988), they represent life-size evolutionary experiments for the study of the mechanisms and consequences of hybridisation (Touchard et al., 2022; Viard et al., 2020).

One particularly interesting outcome of human-mediated hybridisation is adaptive introgression. This process occurs when the incorporation of beneficial genes through introgressive hybridisation leads to an increased frequency of these genes in the recipient populations (Edelman & Mallet, 2021; Hedrick, 2013; McFarlane & Pemberton, 2019; Ottenburghs, 2021). Cases of adaptive introgression have been reported where specific introgressed genomic regions are under selection in association with adaptive traits such as pollution or pesticide resistance (Oziolor et al., 2019; Suarez-Gonzalez et al., 2016; Valencia-Montoya et al., 2020; Le Corre et al., 2020). Adaptive introgression can be viewed as a form of selection on standing genetic variation fuelled by gene flow rather than mutation (Welch and Jiggins, 2014) that allows for rapid adaptation to changing environmental conditions (Norris et al., 2015, Oziolor et al., 2019). Examining changes in the frequency and the spatial distribution of introgressed alleles over time is particularly helpful to understand the conditions under which fitness costs driven by incompatibilities are overcome, the relative role played by selection and gene flow in maintaining locally adaptive genes, and the selection processes at play (e.g., local adaptation vs. strict positive selection). To date, however, there has been little monitoring of adaptive introgression over time (but see Norris et al. (2015) for a detailed study of adaptive introgression to pesticides in mosquitoes).

Anthropogenic hybridisation between two tunicate species of the genus *Ciona*, following the introduction of *C. robusta* in the European native range of *C. intestinalis* in the early 2000s (Bouchemousse et al., 2016a), has resulted in an adaptive introgression island (Fraïsse et al. 2022). The two tunicate species are found in syntopy in ports of the English Channel and the North-East Atlantic Ocean (Bouchemousse et al., 2016b; Nydam and Harrison, 2010). Their life cycles overlap and hybridisation is possible, even though extremely rare in natural populations (Bouchemousse et al., 2016c; Bouchemousse et al., 2017). Crosses in the laboratory showed that there is partial and asymmetrical reproductive isolation between these highly divergent species (Bouchemousse et al., 2016b; Malfant et al., 2018). Despite little evidence showing contemporary hybridization in the wild, introgression by *C. robusta* alleles was found in *C. intestinalis* populations sampled in 2012 in the European contact zone (Le Moan et al., 2021). Introgression is mostly found in a unique and localised genomic region between 700 kb and 1.5 Mb of chromosome 5 and is characterised by long tracts (30-150 Kb) of *C. robusta* ancestry (Fraïsse et al., 2022). This introgression event is recent (estimated about 75 years ago) and not observed in populations outside the contact zone (Fraïsse et al., 2022; Le Moan et al., 2021). Although the introgressed haplotypes are not fixed in any study populations, several pieces of evidence point towards adaptive introgression. Using whole-genome sequences phased by transmission in parent-offspring trios, Fraïsse et al. (2022) found that positive selection on the chromosome 5 introgression island is supported by long-range linkage disequilibrium patterns, haplotype-based tests and the presence of a tandem repeat of a cytochrome P450 gene in the core of the introgression island. This tandem repeat is a likely candidate for selection, possibly linked to detoxification functions often associated with cytochrome P450 genes (Fraïsse et al., 2022).

To get further insights about how selective processes shape introgression patterns (e.g., frequency, tracts length), we examined the spatio-temporal dynamics of introgressed *C. robusta* gene frequencies. We analysed 1,214 *Ciona* individuals, including 1,105 *C. intestinalis* collected during two time periods, spanning about 20 generations, in 20 localities in the contact zone (14 of them sampled twice), from the North Sea to the Bay of Biscay. Because more than a thousand individuals were to be analysed, it was necessary to use a cost-effective genotyping method. Taking advantage of previous genomic resources (Le Moan et al. 2021; Fraïsse et al., 2022) and published genomes (Satou et al., 2019), we developed a KASP genotyping approach, based on 23 ancestry-informative SNP markers specifically designed to distinguish native, non-native and hybrid individuals as well as to identify alien alleles across the introgression island on chromosome 5. We first confirmed ongoing hybridization between the two species is very rare and updated the spatial distribution of *C. robusta* alleles that were detected in new locations for both time periods. Our study also exemplified the benefits of spatio-temporal analyses of adaptive introgression. While we observed stability in both time and space of the frequency of *C. robusta* alleles at the core of the introgression island, we detected decreasing frequencies in the flanking regions (“shoulders” *sensu* Schrider et al., 2015) and weaker linkage disequilibria in almost all populations, demonstrating an erosion of alien ancestry in the shoulders of the adaptive introgression island. Altogether, the observed pattern suggests a kind of balancing selection at the adaptive locus amid counter-selection of alien hitchhiker alleles in the native genome or their replacement by native alleles through gene flow (i.e. the second phase of a local sweep; Bierne, 2013).

## 2. Material and methods

### 2.1. Sampling and DNA extraction

*Ciona intestinalis* and *C. robusta* adult individuals were sampled in 22 ports from the North Sea to the Western Mediterranean Sea during late summer 2021, except for one locality (Dunkerque, site 5) sampled during summer 2022, totalising 775 individuals (Fig. 1, Table S1) (time period referred as 2021 hereafter). More precisely, *C. intestinalis* was sampled in 20 ports from the North Sea to the Bay of Biscay totalising 666 individuals, and *C. robusta* was sampled in four locations totalising 109 individuals. This latter species was easily sampled in two locations nearby the Sète city, in the Western Mediterranean Sea (104 individuals; sites 21 & 22) but although both species were actively looked for in the field along the other coasts of France, only five *C. robusta* individuals could be collected in two of the ports, one in the North Sea and one in the Bay of Biscay (St-Vaast, site 6 and Arcachon, site 19). Species were identified in the field based on morphological criteria (Sato et al., 2012) and a piece of branchial basket was collected and stored in RNAlater (samples from France) or absolute ethanol (samples from UK) for further molecular work. For DNA extraction, when preserved in RNAlater, the tissue was first rinsed in PBS (1X) for one minute, then for all tissues, around 20 mg was digested in 200 µL of lysis buffer (NucleoMag™ Tissue from Macherey-Nagel) and 25 µL of proteinase K at 56°C, 450 RPM for 3 hours in Eppendorf ThermoMixer® C. Extraction was done using Thermo Scientific™ KingFisher™ Flex System and followed the standard protocol of the NucleoMag™ Tissue kit except with 2x diluted magnetic beads. DNA concentration was measured with a Nanodrop spectrophotometer. DNA was normalised at 10 ng/µL using an NxP Span8 protocol.

**Fig. 1:**
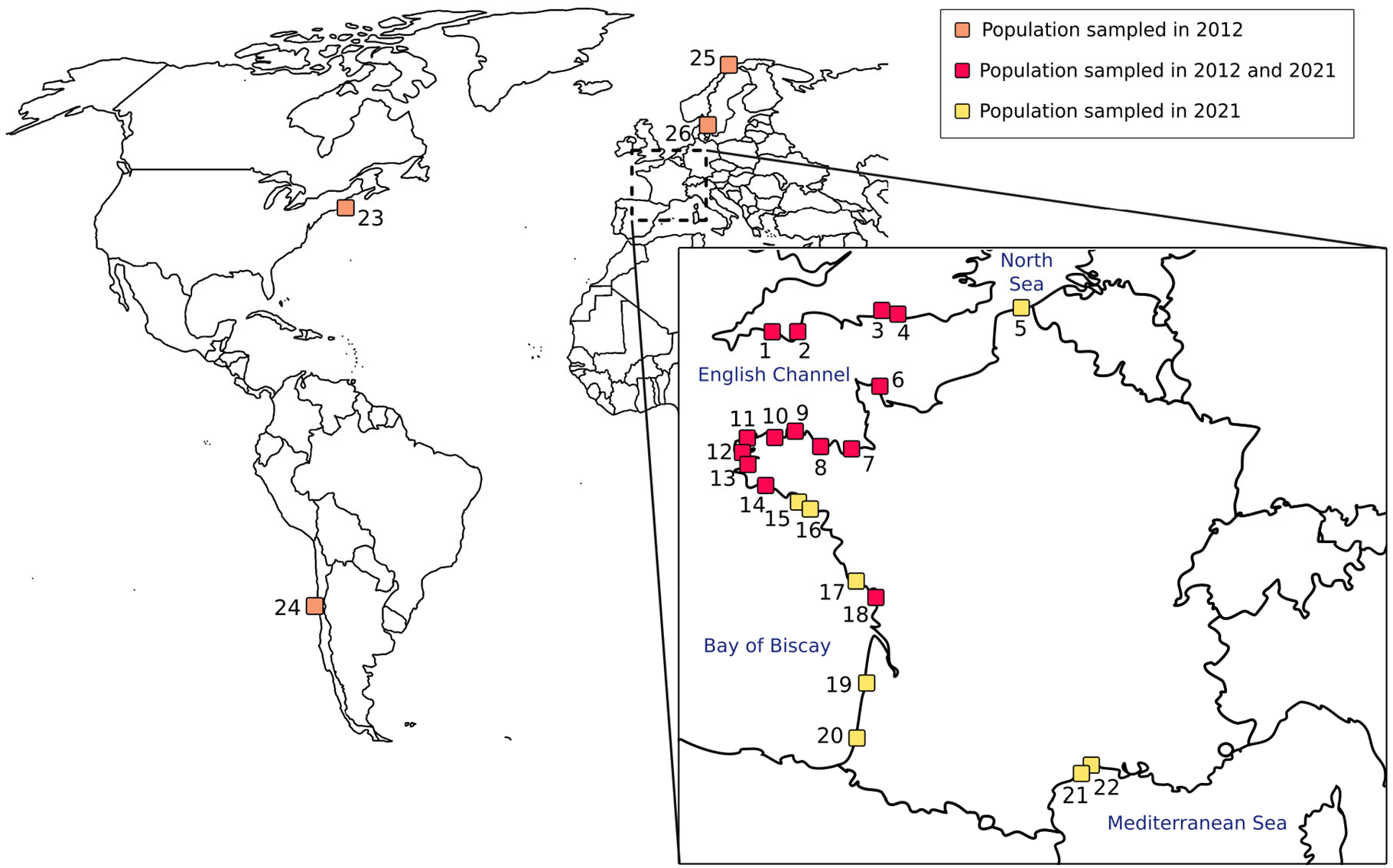
Sampling locations of *Ciona* spp. populations, in the European contact zone along the coasts of France and United-Kingdom (yellow and red squares), and outside the contact zone (orange squares). Locations associated with the numbers on the map can be found in Table S1.

Additionally, for temporal analyses, DNAs of 439 *C. intestinalis* individuals, collected in summer 2011 and 2012 in 14 English and French ports, except for La Rochelle (site 18) sampled in 2014, were included (time period referred as 2012 hereafter). Each of these 14 ports are part of the 2021 sampling (Table S1). For these locations, we included individuals analysed in the previous population genomics study that first revealed the introgression island (Le Moan et al. 2021) and additional ones from the same DNA collection to reach ca. 30 individuals, as for the 2021 sampling.

Finally, DNAs obtained from samples collected outside the contact zone and examined in previous studies (Bouchemousse et al. 2016, Le Moan et al. 2021) as well as nine F1-hybrids were used as controls for our genotyping procedure. Samples outside the contact zone were collected in Coquimbo (Chile; site 24) for *C. robusta* and in Nahant (USA; site 23), Tromso (Norway; site 25) and Gullmar Fjord (Sweden; site 26) for *C. intestinalis* (17 individuals for each location). For the F1 hybrids, one was identified by Bouchemousse et al. (2016c) in one natural population collected in 2012 and eight were obtained from lab crosses using individuals collected in Brest (site 12) and Aber Wrac’h (site 11). These crosses were done similarly and at the same time as the parents-offsprings trios used in Fraïsse et al. (2022).

### 2.2. KASP genotyping

For KASP genotyping, one mitochondrial and 26 nuclear candidate SNPs, each with alleles diagnostic of the two species, were identified from sequences obtained in previous analyses (one from RAD-Seq data, Le Moan et al., 2021; five from transcriptome data, Bouchemousse et al., 2016c; 20 from whole genome sequences data, Fraïsse et al., 2022). The 27 candidate SNPs include i) one SNP located on the mitochondrial genome to keep track of the maternal lineage, as previous studies showed that the hybridisation between the two species is asymmetric, with only *C. intestinalis* being introgressed by *C. robusta* (Bouchemousse et al., 2016; Malfant et al., 2018, Le Moan et al. 2021); this mitochondrial marker allowed to confirm the species identification made in the field based on morphological criteria, ii) 13 SNPs distributed across the genome, one in each chromosome but chromosome 5, to be used in a multiplex with primers pooled together to compute an hybrid index defined as the proportion of *C. intestinalis* allele fluorescence over the 13 SNPs (see Hammel et al. (2023) for a detailed description of this approach) and, iii) 13 SNPs along chromosome 5 to examine both the frequency of introgressed alleles and the extent of the introgression including 12 SNPs along the introgression island and one outside (SNP “SB” in Fig. 2; from transcriptome data from Bouchemousse et al. (2016c)). Using a combination of results from Fraïsse et al. (2022) with genome sequences and Le Moan et al. (2021) with ddRADseq but with larger sample sizes, we identified a SNP shared by the two datasets (SNP 15 in Fig. 2; SNP 38 radtag 877292-877271 in ddRADseq) that maps to the very core position of the introgression island. It is located 311 bp away from the end of the candidate cytochrome P450 tandem repeat. We thus expect a higher frequency of *C. robusta* alleles around this SNP in the receiving *C. intestinalis* populations. We used SNP 15 as a marker of introgression frequency, while the other 12 markers were used to study the extent of hitchhiking on the chromosome.

Primers for these 27 candidate SNPs were designed and produced by LGC Biosearch Technologies. The flanking sequences of each candidate SNP were carefully checked beforehand in order to limit the amount of polymorphism on each side of the targeted SNP and to facilitate the primer design. The GC content in primers averaged 42.8% with a minimum of 18.2% and a maximum of 65%. Primers were tested by genotyping 50 individuals for each SNP. Fourteen DNAs from previous studies with known genotypes (five *C. robusta*, two non-introgressed *C. intestinalis*, seven introgressed *C. intestinalis* and nine F1 hybrids) were used as positive controls. Thirty-six newly extracted DNAs were analysed in the same run. For this testing procedure, one µL of assay mix (KASP-TF V4.0 2X Master Mix, 1X; primers, 1µM; HyClone™ HyPure water) and 0.5 µL of DNA were mixed in qPCR 384-well plates using Labcyte Echo525. End-point PCR was then performed according to the protocol provided in Table S2. The multiplex made of the 13 SNPs mentioned above was tested under the same protocol with all primers mixed together in equal proportions to maintain a target concentration of 1µM.

For each DNA and assay mix, two fluorescences values were measured during the reading step, each characterised by a different fluorochrome colour and associated to a given diagnostic allele. The signal given by the intensity of the fluorescence thus indicates the genotype of a sample at a given SNP. The sum of the two fluorescences was calculated and compared to the value obtained for the negative control to check for bad amplification. The fluorescence data were then analysed using the allele dosage or Variant Allele Fluorescent Fraction (VAFF) which is calculated as 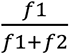 with *fi* the fluorescence specific to the *i* allele (Hammel et al., 2022). The VAFF value indicates if, at a given SNP, an individual is homozygous (high or low value depending on the allele) or heterozygous (intermediate value).

For genotyping accuracy, if the sum of fluorescences was equivalent to or lower than the negative control at a SNP for a majority of samples without any improvement after the recycling step (Table S2) or if the VAFF analysis gave unexpected results for the control samples, the SNP was discarded. To assess the accuracy of the multiplex, the VAFF calculated over all SNPs pooled together, used as a proxy for the hybrid index, was compared to the mean value of all VAFF obtained for each SNPs analysed separately. The correlation between these values was calculated considering two genotyping test runs, the first one with the 50 selected individuals from both species mentioned above (R = 1; p < 2.2e-16; Fig. S1), and the second one with four positive controls from the first run as well as F1 hybrids (R = 0.99; p < 4.4e-11; Fig. S1).

Following the testing procedure, a total of 23 SNPs were kept, including 11 pooled in one multiplex, 11 distributed on chromosome 5 (10 in the introgression island and 1 outside) to be analysed separately, and one located on the mitochondrial genome (Fig. 2; Table S3). Among the four that were discarded, amplification was not working for two of them and only one allele was amplified for the two others. Ultimately, a total of 13 assays (i.e. end-point PCRs) was made for each DNA.

All samples were then analysed according to the protocol described above. For three assays (SNP 20; SNP 21; mitochondrial marker), DMSO was added to the assay mix (5% of the total volume). The recycling step used in the testing protocol did not improve the results and so, additional cycles were not done in subsequent reactions. Four samples (one *C. robusta*, one non-introgressed *C. intestinalis* and two introgressed *C. intestinalis* covering the length of the introgression island) were selected from the testing stage to be used as positive controls in every qPCR run.

**Fig. 2:**
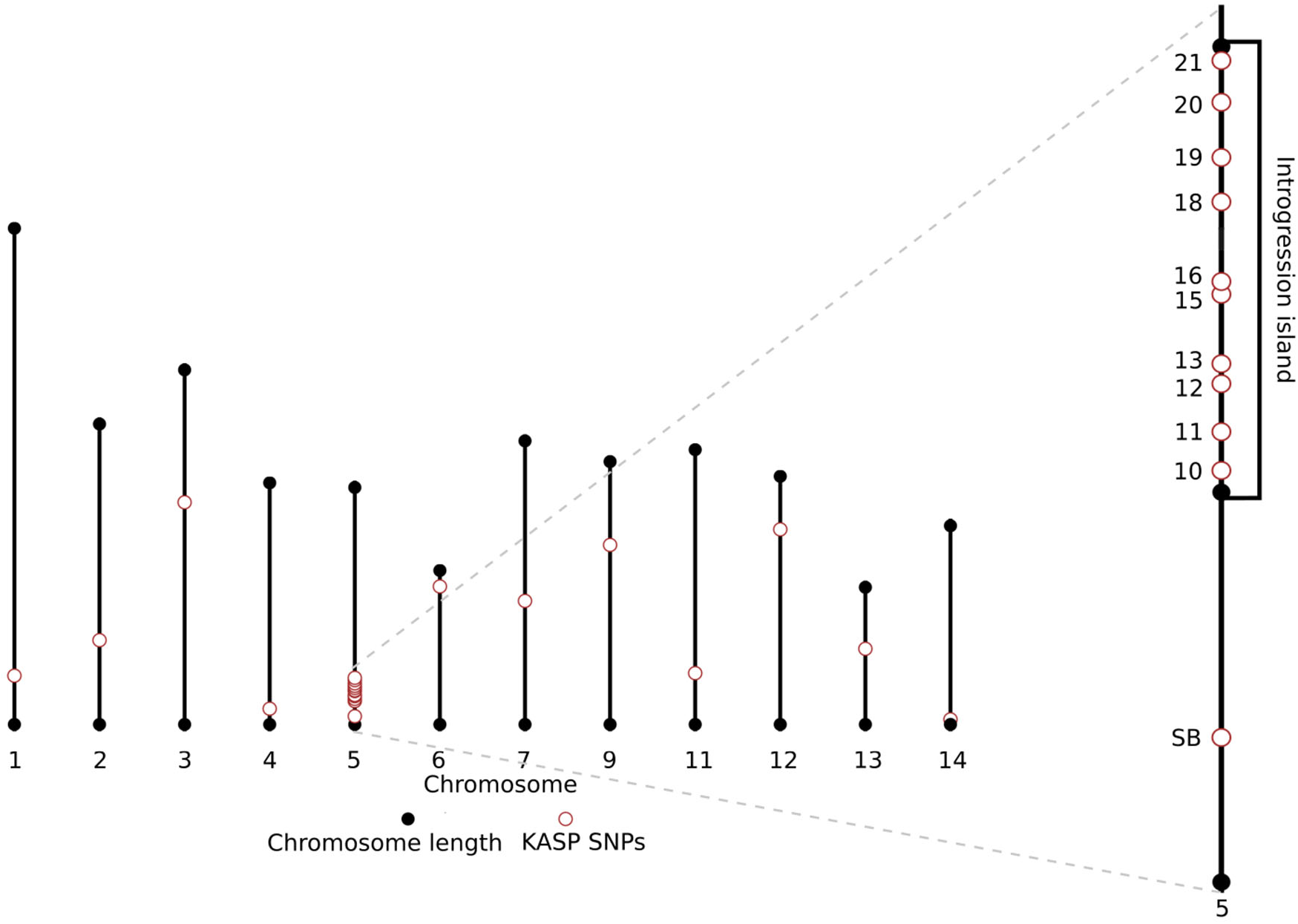
Position of the 23 nuclear SNPs retained for routine genotyping along the chromosomes of *C. intestinalis*, with a focus in the introgression island on chromosome 5.

### 2.3. Data analysis

All analyses were performed in R (v4.2.2). After filtering for bad amplifications, the VAFF was calculated. Based on the VAFF value and the positive controls, three genotypes (homozygous for the *C. intestinalis* allele, homozygous for the *C. robusta* allele and heterozygous) could be observed (Fig. S2; Fig. S3) and were manually assigned to each sample for each SNP by determining cut-offs for each run (Table S4). All genotypes for the introgression island for populations sampled during both time periods can be found in Fig. S3 which allows the visualisation of the introgression tracts.

To determine the presence of hybrids in new samples, the multiplex was first used to compute the hybrid index and assigned each individual to three categories: pure *C. robusta*, pure *C. intestinalis* or admixed individuals (e.g. first-generation hybrids, backcrosses or later-generation hybrids).

The next analyses were performed on a dataset consisting of only *C. intestinalis* individuals, both introgressed and non-introgressed. We first examined SNP 15, at the core of the introgression island, and computed the *C. robusta* allele counts, genotype counts and allele frequency for the two sampling periods. The temporal variation of alien allele and genotype counts between each sampling period was analysed over the whole dataset and per population, by performing an exact test (Raymond & Rousset, 1995) using the *test-diff* function of the genepop v 1.1.7 R library. For each SNP located in the flanking regions of SNP 15 along the introgression island of chromosome 5, the same Fisher exact test was used over the whole dataset. In addition, to examine the directionality of change over time, variations of allele frequencies between 2012 and 2021 were also analysed using a one-sided Wilcoxon signed-rank test using populations as replicates (Wilcoxon, 1992). For combining results across multiple tests, we used the Fisher’s combined probability test with the poolr R library. Finally, to examine linkage disequilibrium between each pair of SNPs and thus infer if introgression tracts are getting eroded, we calculated the correlation coefficient r² using PLINK (v1.90b6.26) across all individuals for each sampling period.

## 3. Results

### 3.1. Genotyping accuracy and verification of species status

Across the 1,291 analysed individuals of both species (including hybrids), only two were removed because of bad amplifications. Missing data after removing these two individuals was less than 1%.

The mitochondrial SNP and the multiplex confirmed the species identification that was done in the field. The introduced species *C. robusta*, which was present in four ports of the contact zone in 2012 (Bouchemousse et al., 2016c), was only found in two in 2021, namely St-Vaast (site 6) and Arcachon (site 19), the former being the only one common to the two time periods. Moreover, the multiplex did not reveal any individual with intermediate VAFF values indicating the absence of first-generation hybrids in the contact zone, for both time periods. None of the 109 *C. robusta* individuals displayed evidence of admixture. Outside the contact zone, two *C. intestinalis* individuals from Norway (site 25) were heterozygous at SNP 13, otherwise no individual was introgressed or admixed in any of the four locations. As Norwegian *C. intestinalis* are outside the contact zone and not introgressed, this is likely due to shared polymorphism between *C. intestinalis* and *C. robusta* at SNP13 that could result in an overestimation of *C. robusta* allele frequencies.

### 3.2. Stable frequencies at SNP15, the core of the introgression island

As expected, *C. robusta* allele frequency was the highest at SNP 15 (1,055,311 bp) both in 2012 and 2021 with an average value of 0.28 and 0.25, respectively, when computing the frequency across all individuals. Allelic distribution was not different between the two time periods (Fisher’s exact test, p = 0.49).

**Fig. 3:**
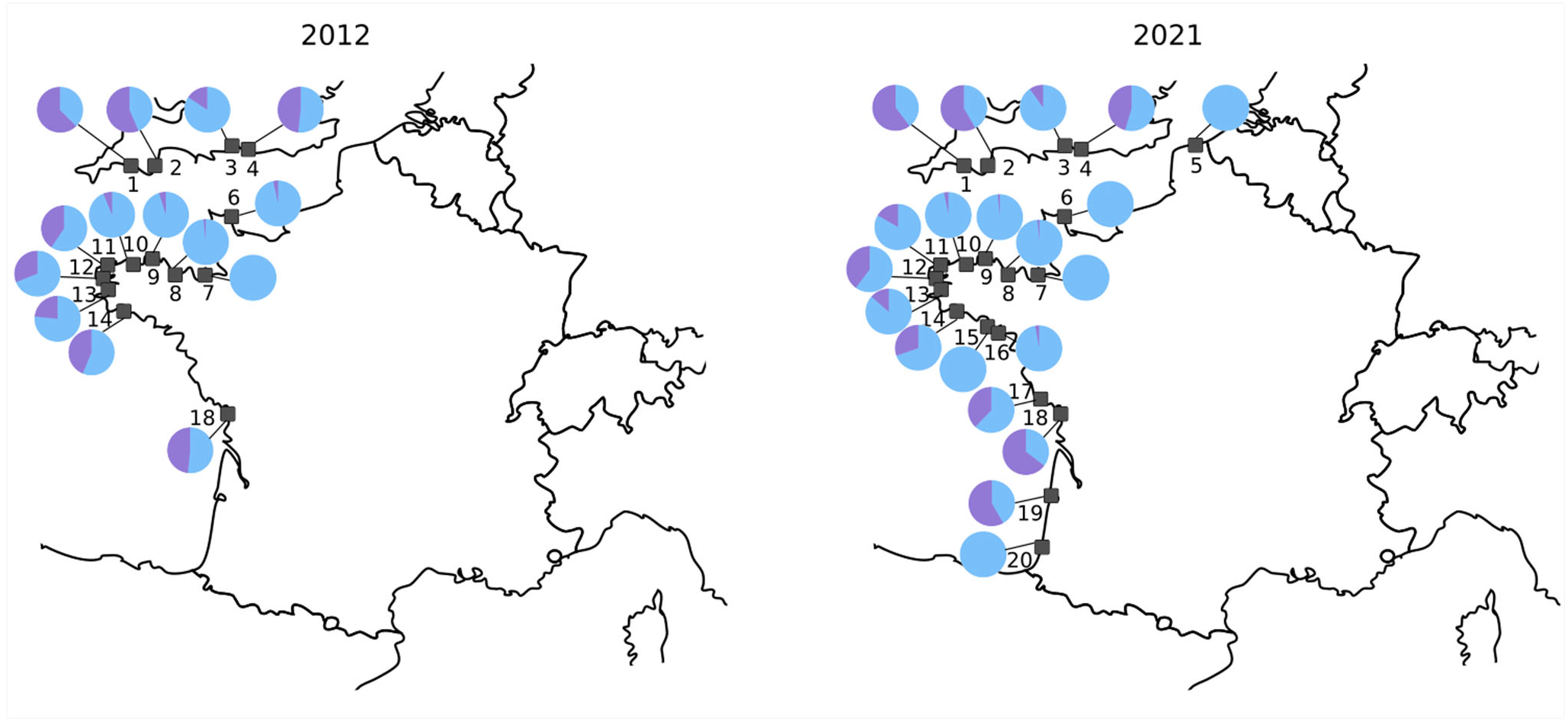
Relative proportion of introgressed *C. robusta* alleles (in purple on the pie charts) and *C. intestinalis* alleles (in blue) at core SNP 15, in 2012 and in 2021, in each study population.

For the 434 *C. intestinalis* sampled in 2012 in the contact zone, 42% (N=183) were either homozygous or heterozygous for the *C. robusta* allele at this SNP. In 2021, out of the 664 *C. intestinalis* individuals, this proportion was 37% (N=247). Genotypic distribution did not show significant differences over time across the whole dataset (Fisher exact test; p = 0.55).

Introgressed alleles at SNP 15 were detected in 13 of the 14 populations sampled in 2012 and in 15 out of the 20 populations sampled in 2021 (Fig. 3). In St-Vaast (site 6), they were detected in 2012 but at very low frequency (0.034) and no longer detected in 2021. Note that in Hossegor (site 20), a newly study site in 2021, only one individual of *C. intestinalis* could be found in the field. This unique individual did not show introgressed alleles at SNP 15, and this sample was not considered in further analyses. For the five other newly sampled populations, two did not show *C. robusta* alleles at SNP 15 (Dunkerque and Etel; sites 5 and 15, respectively) but introgression was detected for the last three (Trinité-sur-Mer, Bourgenay and Arcachon; sites 16, 17, 19, respectively) with an allele frequency of 0.03, 0.38 and 0.58, respectively. When computing Fisher’s exact tests on allele counts within each population separately, there was no significant difference of the alien allele frequency through time (Table S5-A), except for La Rochelle (Fisher exact test, p = 0.049) due to an increase of the *C. robusta* allele frequency over time.

Over the whole sampling zone, we can distinguish three areas where there is a high proportion of *C. robusta* alleles at SNP 15 in the populations at both time periods (Fig. 3): South England (sites 1 to 4), West Brittany (sites 11 to 14) as well as around the Gironde estuary (sites 17 to 19), with *C. robusta* allele frequency at SNP 15 reaching 0.44, 0.26 and 0.55, respectively, when computed over all individuals within these three areas.

### 3.3. Temporal erosion in C. robusta ancestry in the shoulders of the introgression island

Following the analysis of allele frequencies at SNP 15, we then examined allele frequencies at SNPs flanking the core region in the shoulders of the introgression island. The *C. robusta* allele frequency is decreasing with the distance from SNP 15 as expected, although a slight rebound is observed at SNP 20 and 21 in 2012 (Fig. 4) and is even stronger for South England for both years (Fig. 4B; Fig. S4). The allele frequency was lowest at SNP 10 (0.02; 738,970 bp) and 19 (0.03; 1,301,011 bp) in 2012 and at SNP 19 (0.02) and 10, 20, 21 (all three 0.03) in 2021. These SNPs are located at both ends of the introgression island.

When comparing the two time periods and computing the number of *C. robusta* alleles across all SNPs in the shoulders of the introgression island, the mean *C. robusta* allele frequency in each population varied between 0.005 (St-Malo) and 0.29 (Brixham) in 2012 and between 0.003 (St-Malo) and 0.22 (Brixham) in 2021, indicating decreasing allele frequencies in time.

In contrast to the results for SNP 15, among the nine SNPs located in the shoulders of the introgression island, six (SNP 12, 13, 16, 18, 20 and 21) showed significant changes in *C. robusta* allele frequencies (Fisher’s exact test in Table S5B). In addition, five of them (13, 16, 18, 20, 21) showed a significantly lower alien allele frequency in 2021 as compared with 2012 (Wicoxon-signed rank test; Table S5B) (Fig. 4). Combining results across all SNPs located in the flanking regions also revealed significant differences (p = 3.41 10^-6^ and p = 1.91 10^-6^ for allele counts and allele frequencies respectively).

**Fig. 4:**
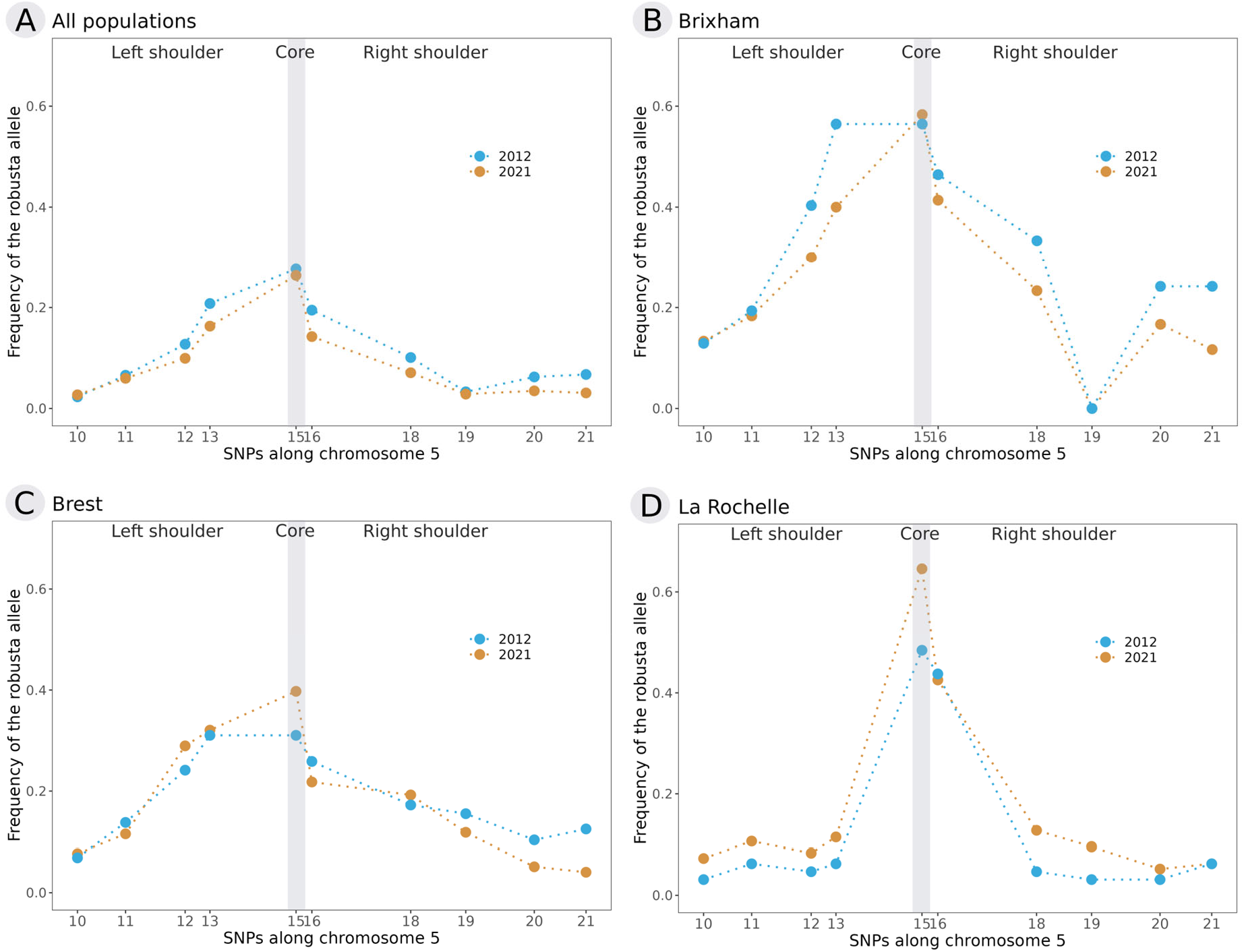
Mean allele frequency in 2012 (blue) and in 2021 (orange) computed at each SNP in the introgression island of chromosome 5 for (A) all individuals (N_2012_ = 434; N_2021_ = 664); and an example population for the three areas with a high proportion of *C. robusta* alleles: (B) Brixham for South England (N_2012_ = 32; N_2021_ = 30); (C) Brest for South-West Brittany (N_2012_ = 29; N_2021_ = 39); (D) La Rochelle (N_2012_ = 32; N_2021_ = 48). The results for the other study populations are provided in Figure S4.

Because recombination is breaking associations between SNPs with time, we expected to observe changes in linkage disequilibrium along the introgression block, notably between the core and shoulder SNPs. The values of r² measuring linkage disequilibrium ranged from 0 to 0.767 with a mean of 0.176 in 2012 and from 0 to 0.642 with a mean of 0.122 in 2021 (Table S6). The most associated loci were SNPs 20 and 21, SNPs 15 and 16 and SNPs 12 and 13. The last two pairs are also the ones with the smallest physical distances on the chromosome (respectively around 22Kb and 35Kb) meanwhile SNPs 20 and 21 are 74Kp apart (Fig. 5A and 5B).

The difference of r² values between the two years was significant, in the direction of SNPs being less associated in 2021 than in 2012 (one-sided Wilcoxon test, p = 0.029), as expected. By comparing values pair by pair between years, we indeed observed a decrease of r² in time for most of the pairs (35 out of 45) ranging from 0.002 to 0.174 (Fig. 5C).

**Fig. 5:**
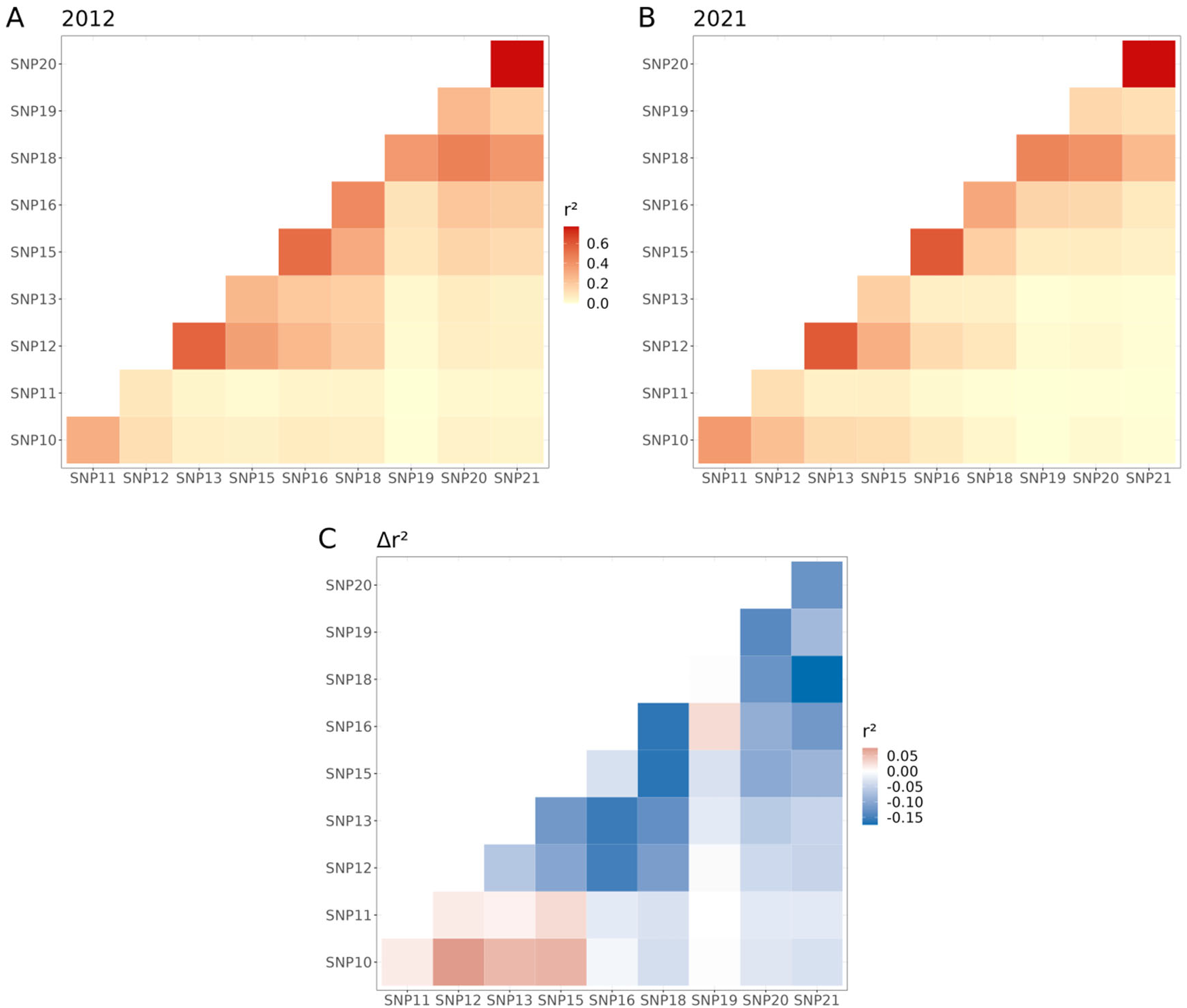
Linkage disequilibrium measured by correlation coefficient r² between each pair of SNPs for both 2012 (A) and 2021 (B). Cells are coloured based on the values, the higher the coefficient is, the darker the cell; r² values are provided in Table S6. (C) represents the difference in r² values between the two years for each pair of SNPs, blue indicating a decrease in linkage from 2012 to 2021, red an increase and white no difference.

We noted above a rebound of *C. robusta* allele in the right shoulder at SNP 20 and 21 (Fig. 4). Linkage disequilibrium values indicate that these two SNPs are more associated to the core SNP 15 than SNP 19, which suggests that they hitchhiked better with the adaptive core than other SNPs. In 2021, the difference of frequency between SNP 20 and 21 and SNP 19 is less important and the r² values are similar between each of these SNPs and the core SNP 15 (r² = 0.060; 0.058; 0.047; Table S6).

## 4. Discussion

We used a set of ancestry-informative markers and a KASP genotyping approach to analyse more than 1,200 individuals of the native European sea-squirt *Ciona intestinalis*, with the aim of monitoring putative changes in spatial distribution and frequency of adaptive alien alleles that had introgressed into its genome, following its recent hybridisation with the introduced species *C. robusta*. Our results show that, in 20 generations, the *C. robusta* allele frequency tends to be globally stable at the core of the introgression island in chromosome 5 while in the flanking regions, we observed an overall decrease of its frequency. Moreover, linkage disequilibrium in this genomic region has decreased in time indicating introgressed alien tracts are getting eroded and tend to be smaller and concentrated at the centre of the island.

### 4.1. Multiplex KASP assay, a cost-effective method for rapid screening of native and non-native Ciona species, and their hybrids

The KASP genotyping approach confirmed the species identification made in the field, with both the mitochondrial SNP and the nuclear multiplex. It also did not reveal any first-generation or later-generation hybrids between *C. intestinalis* and *C. robusta*.

As hybrids between *C. intestinalis* and *C. robusta* were shown to be very rare in natural populations in previous years (Bouchemousse et al., 2016c), the absence of first-generation hybrids in our samples was expected. Our method may lack power to discriminate late-generation hybrids because the multiplex approach is an average of 13 SNPs and *C. robusta* and *C. intestinalis* hybridise very little, but it is powerful to distinguish first-generation hybrids. We thus suggest that the use of our newly developed KASP method, based on the combination of both nuclear multiplex and mitochondrial single-locus assays (simplex), is an effective approach to identify both native and introduced species as well as first-generation hybrids. A similar multiplex assay has also recently been successfully used to infer parental ancestries in *Mytilus* mussels (Hammel et al., 2023).

Regarding the introduced *C. robusta* species, it is noteworthy that this species was found in only two locations St-Vaast (site 10) and Arcachon (site 19) during the sampling made in 2021, despite active search as indicated in the Material & Methods. Interestingly, among the four locations sampled in both 2012 and 2021 and in which *C. robusta* was found in 2012 (i.e., St-Vaast (site 10), Perros-Guirec (site 9), Brest (site 12) and Camaret (site 13); Bouchemousse et al. 2016c)), *C. robusta* was found only in St-Vaast. The KASP genotyping confirmed the morphological-based identification made in the field suggesting that misidentification is unlikely to explain a species-biased sampling. This field observation might be indicative of a decline of the introduced species. It is nonetheless too early to talk about definitive decline and even more of the disappearance of *C. robusta* in this region as this species was reported absent in 2009 except for one individual (in Camaret) but reappeared again in relatively high abundance in 2012 (Bouchemousse et al., 2016b; Nydam & Harrison, 2011). We may, however, hypothesise that introgressed *C. intestinalis* have acquired the ability to now out-compete *C. robusta* in an environment where the latter was previously better adapted. Further monitoring, combining field survey and genotyping based on our KASP multiplex assay, should be done to confirm the declining status of the introduced species.

### 4.2. Stable spatial distribution and frequency of adaptive alien genes at the core of the introgression island

The KASP method allowed us to monitor introgressed alien alleles along chromosome 5. Based on a single KASP assay, targeting SNP 15 at the core of the introgression island, introgressed *C. robusta* alleles have been detected in 13 out of 14 sampled populations in 2012 and 15 out of 20 in 2021 indicating that this introgression is widespread in the contact zone. Our 2021 sampling shows the spatial distribution remains roughly the same and allowed us to detect three new introgressed populations that were not sampled before. These three populations are located North and South of La Rochelle (site 18), which was the only site South of Brittany where the introgression was already studied and reported (Le Moan et al. 2021). Furthermore, re-analysing introgressed populations that were sampled in 2012 with our approach enabled us to screen more individuals than previously done with high-throughput genotyping (i.e., RAD-sequencing, Le Moan et al. 2021). We were able to update the distribution of the introgression reported by Le Moan et al. (2021) for this earlier sampling. Introgressed alleles were indeed detected in two additional populations, St-Vaast (site 6 in Fig. 3) and Roscoff (site 10), where they were previously undetected because the number of study individuals was low. If the introgressed alleles are often rare rather than absent, it means they were given full opportunity to reach the entire study region. This suggests that, if the frequency of these alleles has not increased, they may not be favourable in these populations.

In 2012, the introgression seemed to have a mosaic distribution in the species contact zone (Le Moan et al. 2021), and this distribution is very similar to the one observed in 2021. Three regions are standing out: South England, West Brittany and around the Gironde estuary where *C. robusta* introgressed alleles are much more frequent. It could suggest the initial introgression quickly spread to the other regions thanks to shipping activities connecting these areas. Mosaic distributions are indeed often observed in the case of human-mediated dispersal via shipping (Touchard et al. 2022). In addition, the observed pattern is concordant with the one reported for neutral markers in European populations of *C. intestinalis* (Hudson et al., 2016; Le Moan et al., 2021). A second main observation in our study is that the spatial distribution of *C. robusta* alleles at the core of introgression has remained stable through time. Adaptive alleles can spread quickly in a heterogeneous environment by successive selective sweeps in the habitat where they are positively selected (Schluter & Conte, 2009; Bierne et al., 2013), as observed for instance in sticklebacks where freshwater adapted alleles spread from river to river or in killifish where pollution resistant alleles spread from port to port (Lee & Coop, 2017). We thus have to consider the alternative hypothesis that the observed mosaic structure is rather driven by habitat heterogeneity rather than by the anthropogenic connectivity network of the maritime traffic. Under this hypothesis, populations where *C. robusta* genes are rare or absent, and remained rare or absent in 2021, have environmental components that counter-select alien genes (as is often the case with resistance/cost trade-offs, e.g. Lenormand et al. 1999).

The peak of *C. robusta* allele frequency at SNP 15 (1,055,311 bp) located at the core of the introgression island is consistent with previous studies (Le Moan et al., 2021; Fraïsse et al., 2022). With only a single sampling time, these previous studies were left with the competing alternative hypotheses of (i) global positive selection with the adaptive allele on its way to fixation or (ii) some kind of balancing selection, including local adaptation, that maintains introgressed alien genes at equilibrium frequencies within populations. Our overall observation that alien allele frequencies at the core SNP 15 did not increase between the two periods, spanning approximately 10 years (∼20 generations for this species), supports the second hypothesis. We observed increasing *C. robusta* only in La Rochelle (site 18), suggesting that the selective sweep was not finished in 2012 or that changes in environmental conditions have modified the equilibrium frequency.

Local adaptation is a probable scenario based on our findings. Fraïsse et al. (2022) linked the core region to a tandem repeat of a cytochrome P450 gene (family 2, subfamily U), commonly involved in detoxification processes. The function of this candidate gene, along with the non-fixation and stability of alien genes, indicates that the introgression has now reached an equilibrium between migration (gene flow) and selection. It remains unclear what advantage this introgression provides to explain its persistence in *C. intestinalis* populations. Edelman & Mallet (2021) highlighted that introgression between highly divergent species, such as the *Ciona* species investigated in this study, has to include alleles with strong advantageous effects. These authors linked such a scenario with “extreme anthropogenic selective pressures”, which is also applicable for the two *Ciona* species studied in this research since both inhabit ports, one of which has been introduced. Looking at other cases of human-mediated adaptive introgression similar to ours (e.g. pollution resistance in fish (Oziolor et al., 2019), insecticide resistance in moths (Valencia-Montoya et al., 2020) and mosquitoes (Norris et al. 2015) or herbicide resistance and flowering time in maize (Le Corre et al., 2020)), we can expect that the selective advantage is associated with the specificity of the port’s environment (e.g. pollution, enclosure, substrate, …), where introgressed populations are found. Our diachronic sampling provides further evidence for the hypothesis that adaptive introgressed genes often reach a stable migration-selection equilibrium rather than becoming fixed. The decline in alien allele frequencies in the hitchhiked regions at the shoulders of the adaptive island (see below) also supports this hypothesis.

### 4.3. Falling shoulders ahead: erosion of introgression tracts in the flanking regions surrounding the adaptive core

In the flanking regions on either side of the introgression core (island shoulders), for most SNPs, we observed a decrease of introgressed alleles and a decrease in pairwise linkage disequilibria between the two studied time periods. This is likely explained by recombination breaking association between linked loci after genetic hitchhiking and thus reducing the introgression tract lengths as alien alleles are getting eliminated by purging (Racimo et al., 2015) or gene flow (Bierne, 2010).

The first explanation is that introgression tracts that hitchhiked with the adaptive alien allele carry deleterious mutations in the native genetic background. Given the strong divergence between the two *Ciona* species, we expect most of the alien genome to be incompatible in the native genetic background and to be selected against when introgressed (Schneeman et al., 2020; Schiffman & Ralph, 2018, Dagilis & Matute, 2023). In addition, beneficial alleles must have high selective coefficients to overcome the combined effects of negative linked selection against their flanking neighbours in recipient populations (Edelman & Mallet, 2021). As in Peck’s “ruby in the rubbish” model (Peck, 1994), a beneficial mutation arrives in a genetic background where deleterious alleles are common. We expect recombination in introgression tracts to uncouple this negative selection during the selective sweep, making the hitchhiking footprint narrower than in a standard selective sweep model in which the shoulders are neutral. This expectation is in line with the Dagilis & Matute (2023) model that suggests that short haplotypes are more likely to be introgressed than long haplotypes in divergent species where reproductive incompatibilities have evolved. Once the beneficial allele has fixed or stabilised in frequency, we can thus expect recombination to continue to uncouple negative selection on shorter linked tracts and purge the alien ancestry in the shoulder of an adaptive introgression sweep, and this should be all the stronger as the divergence between species is greater.

The alternative, not mutually exclusive, explanation is that we are witnessing the second phase of a local sweep model (Bierne, 2010). As in a standard selective sweep model, the shoulders of the hitchhiking footprint are assumed neutral. During the first phase, local hitchhiking happens and the locally adapted mutation increases in the population where it is beneficial, bringing with it flanking neutral mutations, all the more so as these are close on the chromosome. Local hitchhiking results in the standard peak associated with a selective sweep, similar to the one we observe here in introgressed *C. intestinalis* populations. Local hitchhiking is fast and is followed by a second phase, interbackground introgression in the shoulders, that can last for a very long time. During this second phase, gene flow between habitats and recombination result in the homogenisation of allele frequencies at neutral loci. This model therefore predicts alien ancestry to be eliminated in the shoulder of the introgression island in populations where alien genes are favoured at the core. It also predicts alien ancestry to reach the shoulders of the introgression island in populations where the core is rare or absent. However, gene flow can be highly asymmetrical depending on demography and genetic determinism, which can explain why we did not obtain evidence for this. At any rate, this prediction will need to be better explored with genome data in the future.

We also noticed a rebound of introgression at the right end of the right shoulder, at SNP 20 and 21, notably in English populations, that was already observed by Le Moan et al. (2021). This pattern highlights the presence of introgression tracts at the right end of the island. A focus on this particular genomic region shows the presence of another cytochrome P450 gene (CYP1F1) at the position of SNP 20. Genes of the CYP450 family often act synergistically on cellular detoxification and clusters of introgression peaks on the same chromosome arm have been found to be associated to the CYP450 family in flies (Svedberg et al., 2021). Another candidate to further examine is the fut8 gene located close to the two SNPs as it has been shown to be involved in glucose metabolism and the cold-response network in *Ciona savignyi* (Huang & Zhan, 2021). However, we observed that *C. robusta* allele frequencies have decreased in 2021 at these two SNPs, similarly to other shoulder SNPs, so we could also propose that this introgression rebound around SNP 20 and 21 would rather be due to chance association during a selective sweep in finite populations, or to negative selection being stronger in the chromosomal region of SNP 19.

Temporal monitoring of selective processes in natural populations is difficult and rarely documented but doing so can offer insights on how it shapes introgression patterns. Here, we show an example of the upholding of alien introgressed alleles in a specific genomic region in some native populations after an event of adaptive introgression followed by human-mediated spread. Meanwhile, hitchhiking introgressed alleles in the surrounding segments are being slowly eliminated by negative selection or flow of native genes, or most likely by their joint action.

## Acknowlegments

The authors are very grateful to Laurent Lévêque for carrying out with FV the 2021 sampling along the coast of France, and to John Bishop for sampling in South England. The authors are thankful to marina operators who allowed conducting sampling in the study ports. The authors are also grateful to Christelle Fraïsse for providing WGS data and advice for SNPs selection and primers’ design and to Alan Le Moan for providing RADseq data. This work benefited from funding through the French National Research Agency (ANR) under the “Investissements d’Avenir” program with the reference ANR-16-IDEX-0006 (i-siteMUSE). FT acknowledges a PhD grant allocated by the same funding. Data used in this work were partly produced through the GenSeq technical facilities of the « Institut des Sciences de l’Evolution de Montpellier » with the support of LabEx CeMEB, an ANR “Investissements d“avenir” program (ANR-10-LABX-04-01). This is publication ISEM 202X-XXX.

## Author contribution statement

FT was responsible for gathering and analysing the data, preparing figures and tables, interpreting the results and writing the first draft. FC was responsible for optimising the KASP protocol, contributed to data acquisition and revised the manuscript. NB was co-responsible for the study design, contributed to the sampling, provided guidance for data analyses, interpreted the results and revised the different versions of the manuscript. FV was co-responsible for the study design, provided funding for the work, organised and carried out the sampling, contributed to statistical data analyses and results’ interpretation, and revised the different versions of the manuscript. All the authors approved the final version and agreed to be accountable for all aspects of the work.

## Data Archiving

The multi-locus genotype table is provided in Zenodo. https://doi.org/10.5281/zenodo.8367591

