## Supplemental Material for "Falling shoulders ahead: adaptive alien genes are maintained amid vanishing introgression footprint in sea squirts"

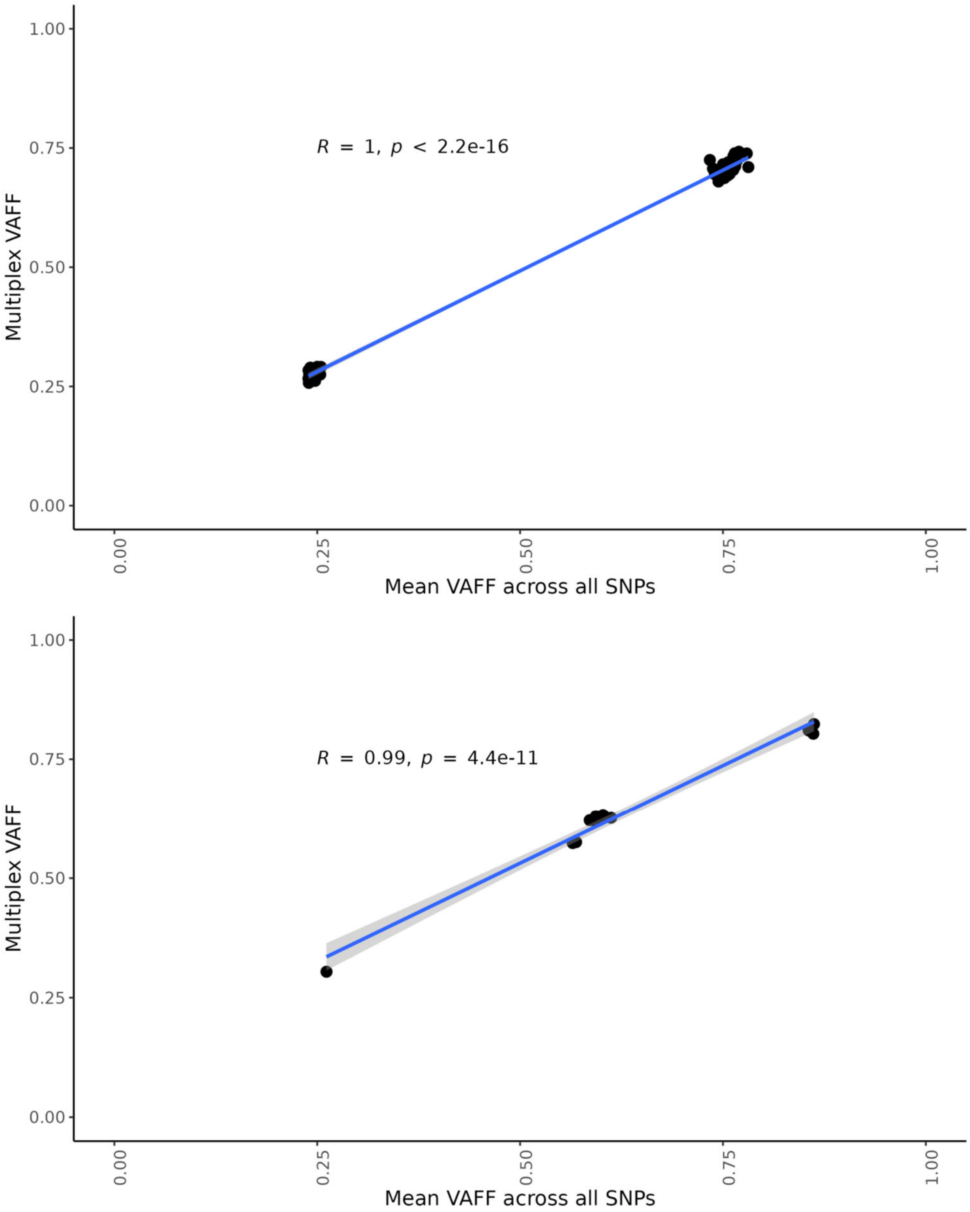

**Fig. S1: Correlation of the multiplex VAFF and the mean VAFF calculated across all SNPs analysed separately (simplex) with, for the top figure, 50 test individuals from both *C. robusta* and *C. intestinalis* and, for the bottom figure, with *C. robusta* and *C. intestinalis* controls chosen from the test individuals as well as F1-hybrid individuals (one found in natural populations in 2012 (Bouchemousse et al. 2016c) and 8 obtained from crosses in the lab produced at the same time than parents-offspring trios used in Fraïsse et al. (2022)).**

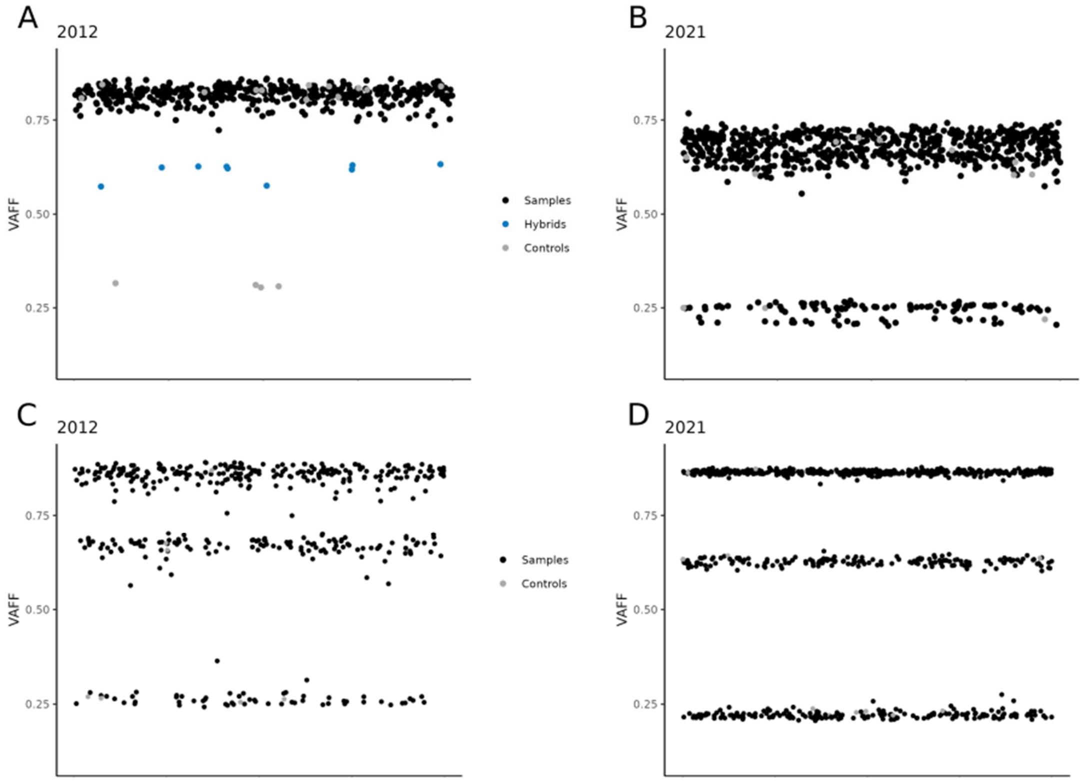

**Fig. S2: Distribution of VAFF values for the multiplex for samples collected in 2012 (A) and 2021 (B) and SNP 15 at the core of the introgression island for samples from 2012 (C) and 2021 (D).** Study individuals (black dots) are shown with controls, i.e. parental species, introgressed and non introgressed *C. intestinalis* (grey dots), and F1-hybrids (blue dots).

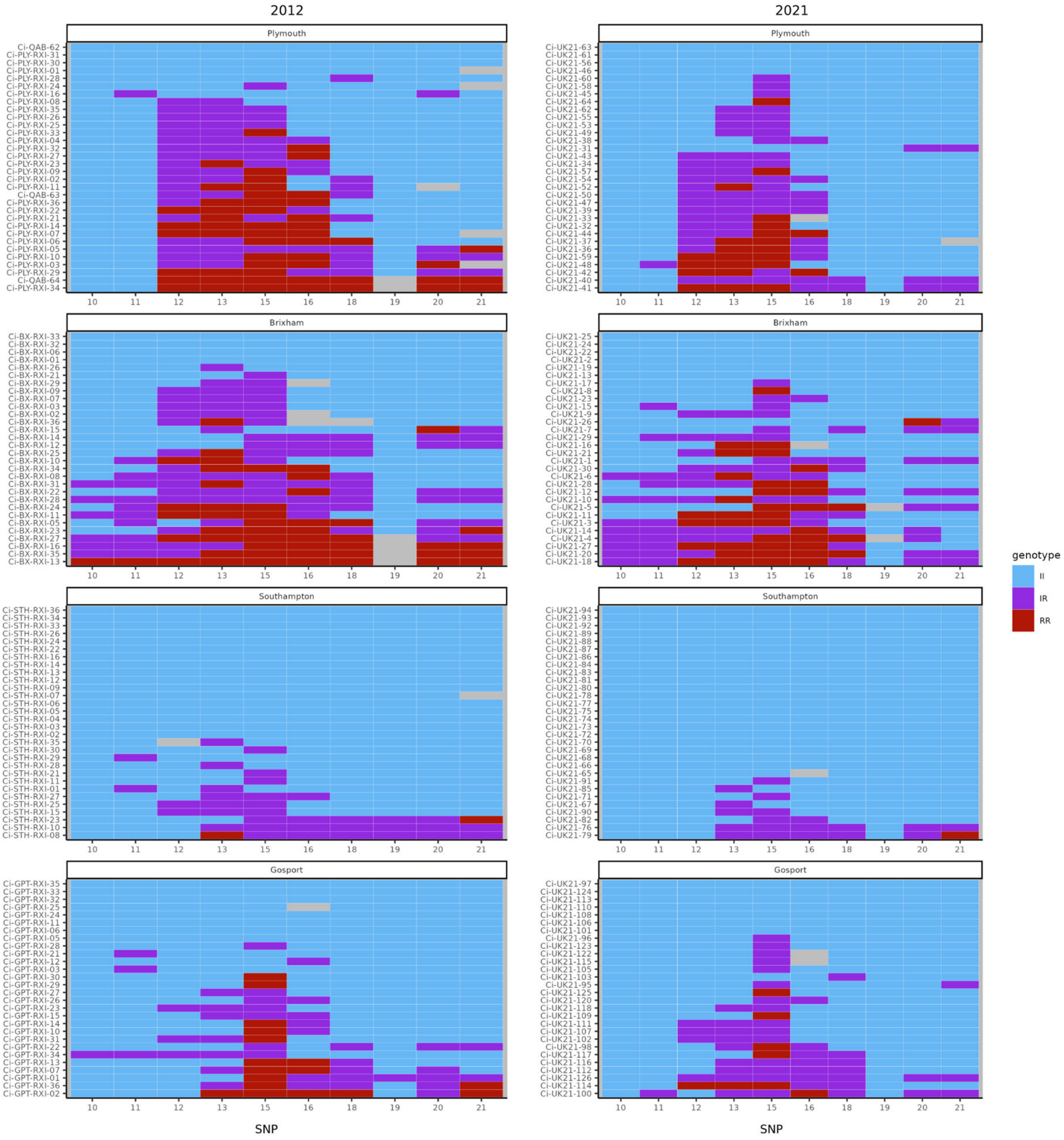

2012
2021
Plymouth
Brixham
Southampton
Gosport
10 11 12 13 15 16 18 19 20 21
SNP
genotype
II
IR
RR

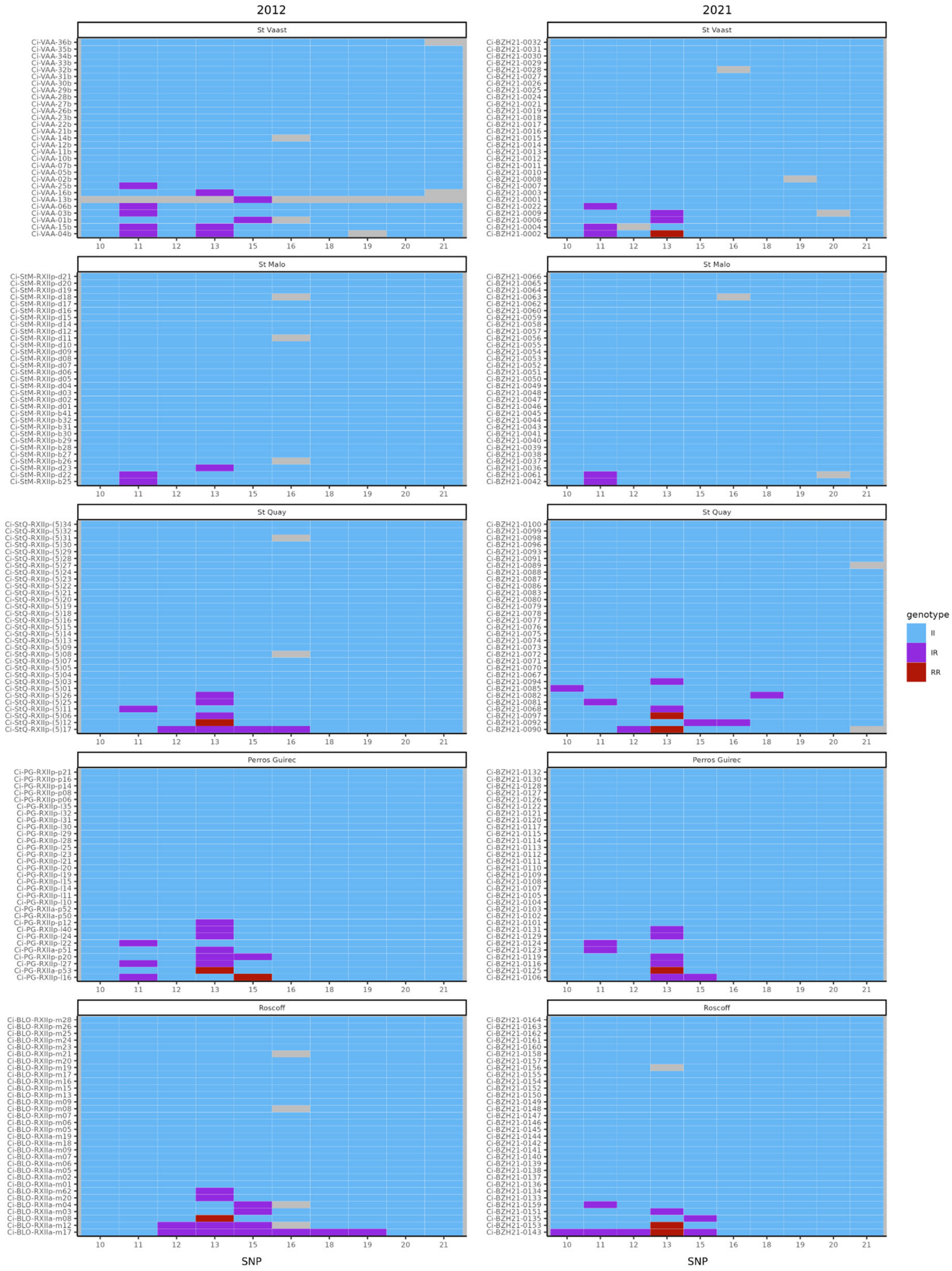
2012
2021
St Vaast
St Malo
St Quay
Perros Guirec
Roscoff
10 11 12 13 15 16 18 19 20 21
SNP
genotype
II
IR
RR

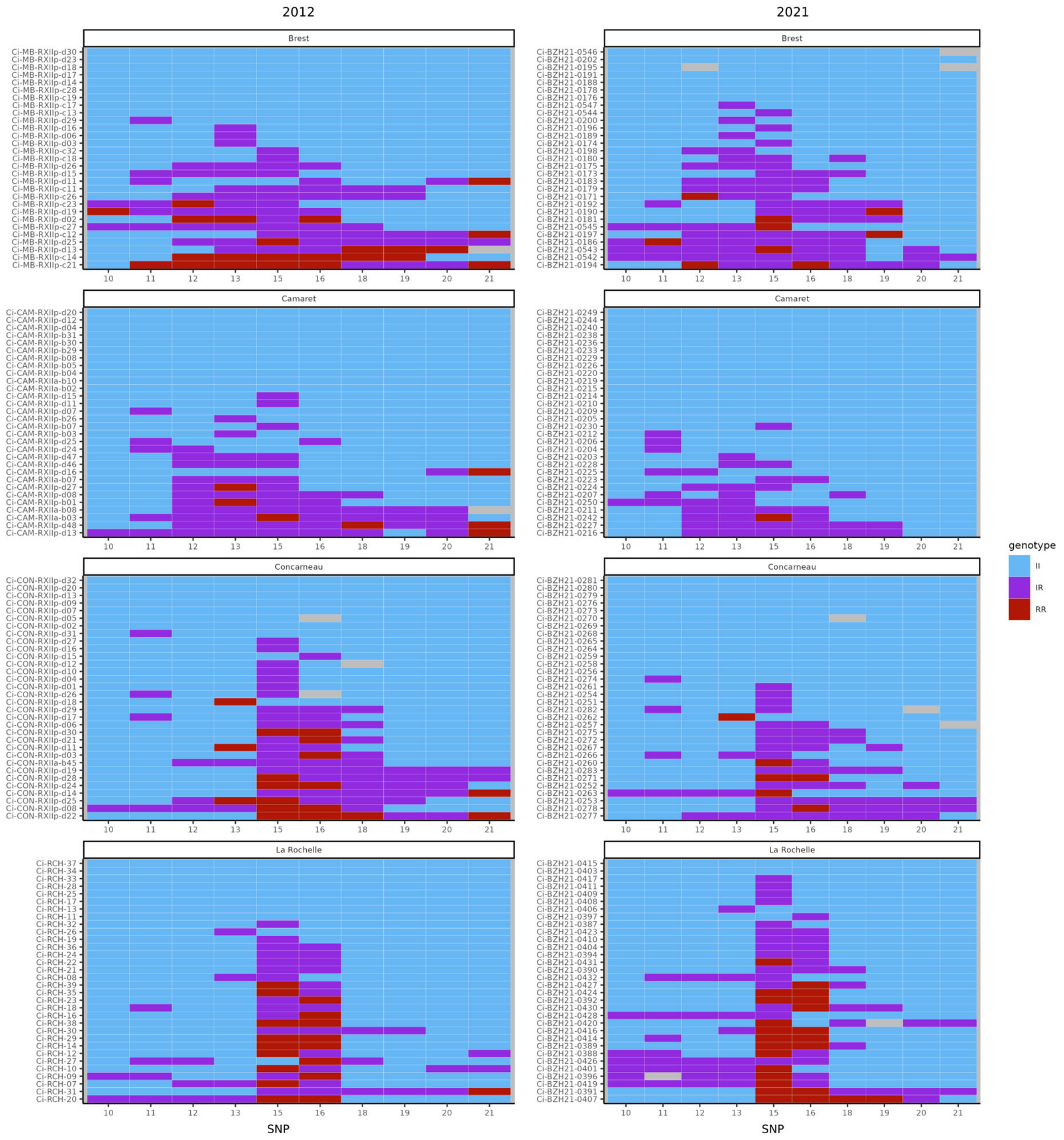

**Fig. S3: Genotype of each individual for each SNP genotyped in the introgression region of chromosome 5.**

For one given population, left and right panels correspond to 2012 and 2021 sampling, respectively. Blue is for *C. intestinalis* homozygotes, red for *C. robusta* homozygotes, purple for heterozygotes and grey is for missing genotypes.

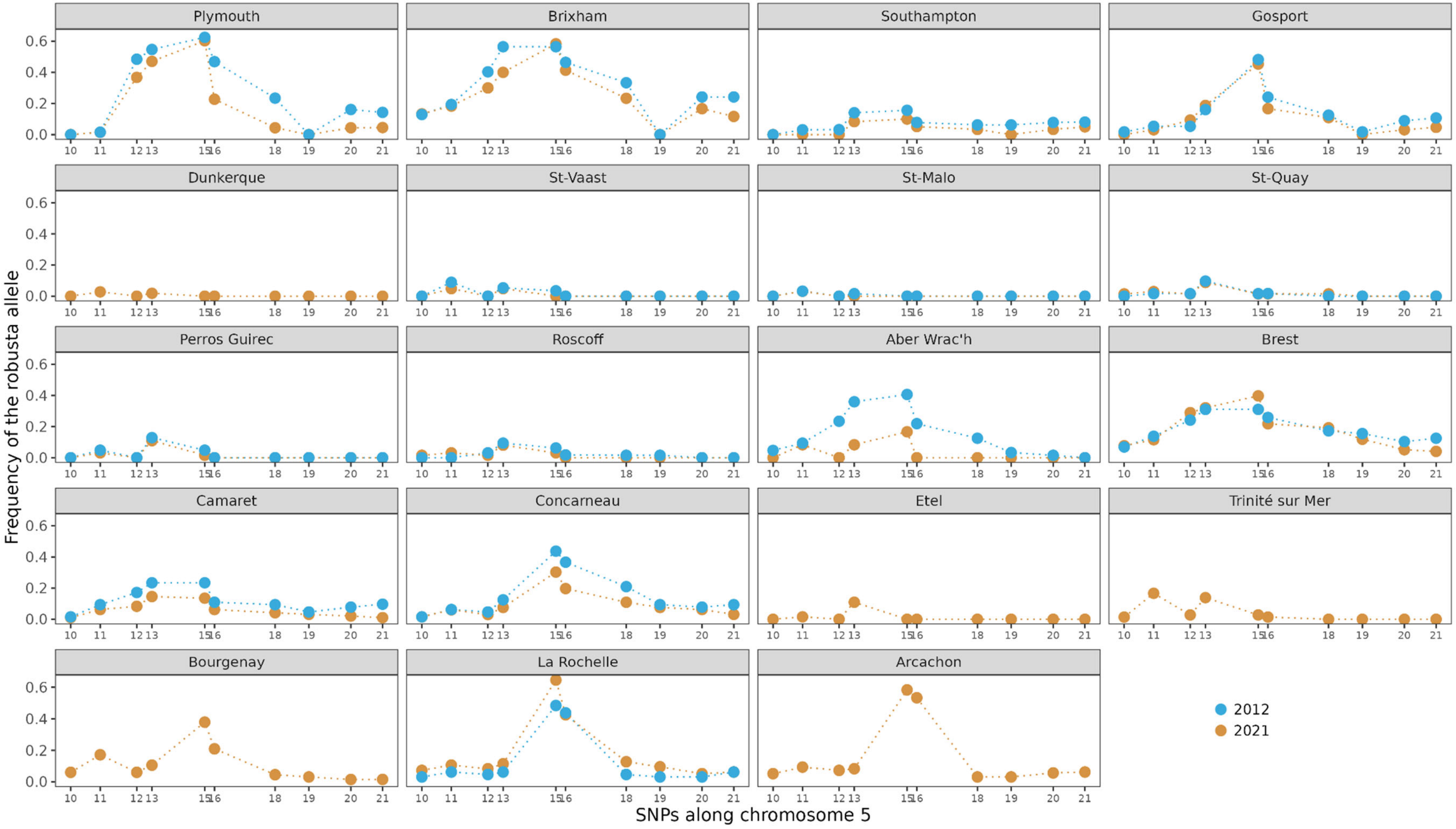

**Fig. S4: Frequency of the introgressed *C. robusta* allele for each SNP analysed along chromosome 5, in each population and for the two sampling periods spanning 20 generations (2012 in orange and 2021 in blue).** Note that the sampling size is balanced across populations (ca. 30 individuals), except at Aber Wrac'h where only 6 individuals could be sampled in 2021.

**Table S1: Sampling number per localities, time period and species**

The number (as shown in Figure 1 in the main text) and name of the sampling location are provided with the number of individuals collected for each time period and species. Ecoregions were named according to the classification of Spalding et al. (2007).

| Location number | Location name | Ecoregion | Number of individuals sampled in 2012 | | Number of individuals sampled in 2021 | |
|---|---|---|---|---|---|---|
| | | | ***Ciona intestinalis*** | ***Ciona robusta*** | ***Ciona intestinalis*** | ***Ciona robusta*** |
| 1 | Plymouth | Celtic Seas | 32 | - | 34 | - |
| 2 | Brixham | Celtic Seas | 32 | - | 30 | - |
| 3 | Southampton | North Sea | 32 | - | 30 | - |
| 4 | Gosport | North Sea | 30 | - | 32 | - |
| 5 | Dunkerque | North Sea | - | - | 54 | - |
| 6 | St-Vaast | North Sea | 32 | - | 32 | 2 |
| 7 | St-Malo | Celtic Seas | 32 | - | 32 | - |
| 8 | St-Quay | Celtic Seas | 31 | - | 34 | - |
| 9 | Perros Guirec | Celtic Seas | 31 | - | 32 | - |
| 10 | Roscoff | Celtic Seas | 32 | - | 32 | - |
| 11 | Aber Wrac'h | Celtic Seas | 32 | - | 6 | - |
| 12 | Brest Moulin Blanc | Celtic Seas | 29 | - | 39 | - |
| 13 | Camaret | Celtic Seas | 30 | - | 48 | - |
| 14 | Concarneau | South European Atlantic Shelf | 32 | - | 33 | - |
| 15 | Etel | South European Atlantic Shelf | - | - | 32 | - |
| 16 | Trinité-sur-Mer | South European Atlantic Shelf | - | - | 36 | - |
| 17 | Bourgenay | South European Atlantic Shelf | - | - | 33 | - |
| 18 | La Rochelle | South European Atlantic Shelf | 32 | - | 48 | - |
| 19 | Arcachon | South European Atlantic Shelf | - | - | 48 | 3 |
| 20 | Hossegor | South European Atlantic Shelf | - | - | 1 | - |
| 21 | Etang de Thau | Western Mediterranean | - | - | - | 32 |
| 22 | Sète | Western Mediterranean | - | - | - | 72 |
| 23 | Nahant | Carolinian | 17 | - | - | - |
| 24 | Coquimbo | Central Chile | - | 17 | - | - |
| 25 | Tromsø | Northern Norway and Finnmark | 17 | - | - | - |
| 26 | Gullmar Fjord | North Sea | 17 | - | - | - |

**Table S2: End-point PCR protocol for KASP genotyping.**

| Step – Number of cycles | Program |
|---|---|
| Initialisation | 15 min at 95°C |
| Amplification 1 – 10 cycles | 20 s at 94°C then 1 min at 61°C |
| Amplification 2 – 29 cycles | 20 s at 94°C then 1 min at 55°C |
| Read 1 | 1 min at 37°C then 1 s at 37°C |
| Recycling – 3 cycles | 20 s at 94°C then 1 min at 55°C |
| Read 2 | 1 min at 37°C then 1 s at 37°C |

**Table S3: Flanking sequences and diagnostic alleles for the 23 SNPs kept for routine genotyping (see main text for details).**

| ID | Sequence | *C. robusta* allele | *C. intestinalis* allele | Position in genome |
|---|---|---|---|---|
| mitochondrial | ACTAGRGTTGATATAGCTATTTTTTCTTTRCATTTAGCTRGRGTTTCTAGTATTTTAAGATCAGTTAATTTYTTAGTTACYTTATTTAATATAAARAATAAAAGAAARTCWATAAGTAAYTTAAGTTTATTTTGTTGATCTTTAATTGTTACWACTATTYTMCTAGTAYTATCWCTTCCWGTTTTAGCTGCWGCWATTACNATATTATTATTTGAYCGWAATTTTAATACTACKTTYTTTGATCCKAAYRGRAGAAGRGATCCWATTTTATATCAACATTTATTTTGATTTTTYAGACATCCAGAAGTTTATATTTTRATYYTACCWAGATTTAGAATAATTAGTCATGTAATTGTYTTTTAYTCYAGAAAAGATAATATYTTTAGGTATTATAGAATRGTTTGAGCWATRAGRGGWATTAGRTTTCTTAGRTTTYTAGTRTG[A/G]GCTCATCATATATTYAGTGTARGWATAGATGTTGATTCTCGAGCTTATTTTACTTCWGCTACNATAATTATTGCWGTTCCTACWAGAATTAARGTRTTTTCMTGAGTTTCAACWTTATTAAGRGCTAAWATTYAYTGARGATTACCWCTWTTATGAGCWTATRGATTTTTATTYTTRTTYACTATTAGARGRTTAACTAGAATTRTTTTAGCTAATTGTAGTTTAGATCTTGTYCTYCATGATACWTATTATGTRGTTGCTCATTTTCATTAT | T | A | |
| chr1 | GGTGAAGGGGGAATAAGYCTAACTGCACAGAAGTGCCCAATATTCCAAG[C/G]CCATGGCACATCAGACCCCATGYTRCCRTTCCAGTTTGGSCAGATGAGYAACATGCTGTTACAGAATGCAAGRAAGGATCTNAATCTHACCACTGAGTTTAAACCTTATCAAGGCATGGGRCACCAGTCATGTGAYG | C | G | 1543504 |
| chr2 | GTGTCTTGGGTATCAAGGTTAAAATCATGCTTCCTTGGGAYCCACAAGGTAAAATCGGACCCAAGAAACCTCTGCCTGACAATGTRAACATTGTTGAACCTAAAGATGAAGAAGCCATCACTGGGCCAAGGAGTGARTCCAAGGTTACCAAGCCRATGCCAGATCCCATGCCTGCAGC[T/C]CCACNGCAAGCAGCTATGCCTCCMATGCAGCAGGTYCCACCCCAGCAAACCAWGCCACCTATG | T | C | 2666317 |
| chr3 | ctttaaaatatacagtttaaaactttmccaCCGCTCAAAAAAACATTGGAGTGCTTTTTCCATTyAATAGCCGCGAACAAACGAkTGATTATGACTCGACAATAGAAAACGCGAAAAACGCrACCAAAACGATTTyGGCGTCACGGTTGA[G/A]CGACGTCACGTGTGTTGACTCAGCATTTCCAyAGCAAATCTTTCGAGAAACGAwGTCGATACATTAATAGTTACAATGTGTTTGTGTGTTGCaatgtgtgtgtttgtgtttgtgtgtCTGTTCTCTCkTGTACGGtcatgtrtttgtttt | G | A | 7030902 |
| chr4 | YATATGTAAGTTTCTATCAAAATGTCAGAA[C/T]AAAGATGGTGGTTTTGGTGGTGGCCCTGGTCANNNNNNTCACCTTGCRCCCACTTATGCTGCAATTAAYTGTCTTTGCTCAATYGNAACCAARGARGCTTACTCYGTCATAAAY | C | T | 497855 |
| chr5_SB | GTACAACCATTTGGTGTGGTGAATAATACATTTGGTCGAGCATTAGATTGTGAAGGTTGGTTYACTATTGGWATATGGACGGGCYTGATTGTCACTTTATTATTGGTAACAATTCTTACTCTTGGACT[A/C]TGCATGATTGCACAAATCACTACAATGGACAGATTTGATGATCCRAAGGGAAAACAACTTTCCATACCYCAACAAGAS | C | A | 259572 |

| | | | | |
|---|---|---|---|---|
| chr6 | taaatagacACCATTGTTGAATCATAGTTCTTTAATTGTGTCCTAACAaatgt<br>tattgtgatgtcatttatCkTGTATAACAGCAACTGTTGtgtgrattmtttttatygtcttaatattt<br>taagttcaTAATGAATTGAATAACTGAtc[A/G]attgttacatcataawcatGAATTGT<br>TTACAGGAGATTAAGCATTGTTACCTCACACTACTTGATCCCTAGTTA<br>GrGGAAACCCCCACCATAAATAATkGCATGAACttttatcccatcttatcatttataat<br>tagttttaaacattttt | A | G | 4367328 |
| chr7 | ATGAGTTCTGCCAACAATGAAAACAAAGCTCC[T/C]GAGAAAGCTTCA<br>ACATCTGGTCGTGCWAMTGCTAGYTCHGCCAAAGATAATGCAAGTG<br>ACACTTGGTCACTTAAAAACTTTGAYATTGGCAAACCAYTRGGACGA<br>GGAAAGTTTGGCAGCGTGTACCTTGCTAGAGARAARAAGAGCAAGT<br>TTATCGTTGCACTGAAAGTGCTGTTTAAATCRCAACTYATGAMRARTA<br>ATGTGGARCATCAATTRCGAAGAGAGATTGAAATTCAGTCTCATTTR<br>CGTCATCCACACATTCTGCGACTTTACGGTTACTTTCACGAKGAGAC<br>GAGAGTGTATYTGATCYTGGARTATGCATCTCGTGGGGAAATGTACA<br>ARGAGCTGCAGAAGCAGGGCAARTTTACAGMGGAGMTGTCCGCYA<br>CGTACATAGCYGAGCTMGCAGATGCRCTCAACTAYTGCCACAGCAA<br>ACARGTCATTCATCGTGACATCAAACCKGARAACTTRTTGATGGGTC<br>TTCGRGGGGARTTRAAGATTGCTGATTTTGGTTGGTCTGTGCATGCT<br>CCGTCYTCTAAACGCCAAACYCTTTGTGGTACRCTTGATTACCTCCC<br>RCCAGAGATGATTGAAGCAAAAGATCATGAYGCTAAYGTYGAYCTGT<br>GGACACTTGGSATTCTATGCTATGAGTTTCTTGTTGGCAAACCACCY<br>TTTGAAACAAAAAGCACACAAGAAACATACCTHAGAATYACATCACT<br>GAAATATKCATTCCCTCCCCATGTATCAGAGGGAGCWCGTGATCTTA<br>TTCGTCGCCTSTTAAAGTTGGAACCRCGACAYCGWCTTMCACTRGA<br>AAGTGTRATGGCTCACCCCYGGATCARAGCCMATGC | T | C | 3909404 |
| chr9 | AGTTCAATATAGAACAATATCGACAAACATGCACAGACGGGRAAAAT<br>CAATASCAGTCTTTGTCGATTAAGTTAATCTCCGCTCTTAATATCCTG<br>TCAAASCGTGTACTTGTTATCCTTTGCTATTKCGCTGTGTTATAATTTA<br>GSGTCA[T/C]CGTTATAKAGCTKRAGGTACGCCRRGTTTATACAACAT<br>CGCACKTTTCCTTGTTAGTAGCTTTGAAARCGATTCTCMACACGTTTT<br>TGGTTCTTAATAATTGATAGCAGCACCATTAAGCCGAMCGTCTGGGC<br>TCACTGAAAGAAATTGT | T | C | 5680810 |
| chr11 | GAAAMNRRAGGADTWCGARGAGATTATTTYATTTRTTAAGTCAAACA<br>AAWGCAAGAAATTAAAKRRNTTRATCWACCAAGTAAAHGATAWGTTT<br>GAAGYYAAATTTTCCTACCAAACATTRGTMAGCATATGTCTCATACAC<br>CACCAAAACCATGTRAAAAGAACWAKTCATGATAAAAGCACARATGY<br>NWRYRTRGAWAAATACTATTCYACMTTTTTACAAAGYAAAAKARATR<br>RAAAARGCCAYRAAAACTCAGCAYTGRTGRAAATTGCAACYGAAWA<br>CRAYATTCCACCTTTAYTRCTNGCTAGRTTRATTTTACATAGACWYCT<br>WAAATTAAAAGCASABVAAAGYRACCATRYKGRAACARVYVAAARAG<br>TTAATYTYAAAACTGARGTYGCTCGACTKGTAAAAGATCCMTTTTTWA<br>TCRAAGACMAAGKTTTMTCATGGGAAGTAMGRCAVTGCATTCTACAT<br>GATTTTGGTTATGGCCAAGTAACTGATTCAGTACGARGTCTYATTGG<br>TTCAGAATATGAACACAAAMTAGTGACGHAYGTGAARVRRTTRAATA<br>TTCCATTTCARACYGARGTTGATTTAAAGAAATTGGGATTTGATAAAA<br>CACCRGATATTAAACTTGAAATCCCATTTAAGTTRAAAVACCAAGTGG<br>TKTGCTGGATTGAAAGYAAAGCTTCTTTBGGCACTCCTGAAGAACAT<br>DTGTATTATGTGAATAAACAATACAA[C/T]AGCTACTGGAATCGATTTG<br>GGCCAGGACTGGTGATCTATTGGTTTGGTTTTGTTRATGARTTGGCG | C | T | 1623797 |

| | | | | |
|---|---|---|---|---|
| | YWWGATATGCTCAAGAAGAATATCCTKGTTATGGATAGATTTCCAMTTCCTGCVGAAKTKTCYTTTTWTAATCCTGCTTTWGTTGTWAAAGATGATAARCTATCA | | | |
| chr12 | CCACTCACGACAGTACCAAAGCGTGTTTCCACTTCGAGGMCCAACGARCACGACCGTCTGCTGCACTATCACTCgacattatKacgtataYaTAGGCGCGGGCGACCGCACGAAAGACGCAACAGAGCGAAATCTWSAGTTCTCGCTGTR[A/G]AACCACAGMTTTGTTACATGCGCTACAACTTTAACGGCGATTTTTACTGTTTAGTTTGCGTGTCYACAGTAGCGCTCGTGTTTCCTGTAAGCGTCCGSCGTGGCTTGTTGCCGTTTCCCGACTRCATGGTCGTGTTGTGTCGTCACGGCG | A | G | 6173939 |
| chr13 | CGGYCGGCCKRTCGGTTAAATAATTCTCGATRTAATTTTCGGCGCATCGGTTACGGCTWGGGGGACGTGGGAGTGTGTTATGTTTTGGTGTYGAYTCTAAATCGGTACGAGGRGGCACTGGRGGGCAGTCRAGRTCATCGCTGGTRATTC[G/T]TTTYAAGCGTAACGTGACGTCACCKCCCATGTCRCTCGCATTCCACGgaaaactgaaaacaaaaaacaatatttaggcataaatttaaWttgtgaaaWaattgAACTGTCGatgtatttatMtcttcgaatacagtagcGAATTCAAACA | G | T | 2394321 |
| chr14 | GMGTCTAATTAAAAAATCGCTTCGTYTTTTTACAAATTGCRTTCAAGTGRCRCAAGAAAGATCAAATATCccaaattcaWttaaataacttaaatacatTTCGAACATTGTAACTGTATTGMGAGGACCACTGCRGCAAACCAGCGCGAT[G/A]GGAATTGCAGCTCTTACRTGGCAGGTGTTTGAACAAGTTGCCACTAAACTTCTAGTTGTTCTCTCTCATSTCTTGGTTTATAATRAGCGAYTCGCCCAAGTATCTGATACGGACYTCACCCAATCCTCTATTGYTTGCTTCTAYTCTGTG | G | A | 159786 |
| chr5_10 | ACCTCCACTAGATTTYAATGTCTAAGWAMTYSCATGTTTGTTAAGGTCAAAGGAGTTTTCAAGYTTCATATCTCGATGTTTGGTRAAAGATCCAACRAAAAGAGCRTCGYTAGCCGAGTTGCTTAACCATCCTTTCCTCACCAAYATCTC[G/A]TCMCCACAACCTCTTGTKAAGTTGATCGCTGAAGCAAAAGCTGATGTCACAGAAGAAGTKGAAGAACCTGTGagaattattttgtYYRttttaaatYtgttGATTWCCAATGTGTTCTGGTTTTATGAAAATTTGAACCTAGTTAAGACT | G | A | 738970 |
| chr5_11 | ATACAATCYCARCYTGTGAACAWTCTTACTCATGAGCAACTACGGGTTGTGAATCATGATTTTCAAACTGGTCAGTTGATWCGAGTAATGGCTTATGCYGGTGCAGGAAAAACAAGCACRTTGATYGCTTGTGTTAAAGCTAAGCCAAAC[A/C]AGACATTCCTATACACCTCATTTTGCAAGTAGGATTGTTTTACAATGTCTATAAATCTTGTYACTRRAAgRttcagttttttttctgttataatttMtgttactgtttttaacaataggCAACCATAAACAATTATTAGTCTGRTAARG | A | C | 808581 |
| chr5_12 | RCARAWAATTGATATTAGTAACCACCGTTCAATCATTACCACCTMGGCTTACWTTCAAATGCGTCATTTGCTTGTTGACGTGACCACAGACTGTCSTCGTASGCRGCGCTTTGAATRATYACATTCGAAAATAATMCTTCGYTCTTTGGA[T/G]AGAGGGCATGCATGTGAACSGAGGAGCCACCGGCCTCTTCRCCAAATAKAGTAATTYKAGYWGGATCGCCGCCGAAMgcctgtaaaaaaWtacaacWYtTAGRTAARCCTATTAAGGACCRCAAACACAgatataRgtttttaaataaWa | T | G | 894811 |
| chr5_13 | WAACATGTATGAGCCTGGAGATGGTATACCCCCACATACYGAYAACACAAGATCYTTTGATGGTGTGTTATCMACYGTAAGCTTGGGTTCACATACTGTTATGAACTTTAGYAAAGAMGGTGCTGAAAGAATAGATGTGTGTGTCGAGCC[G/A]cgtactttatttttatttactggtGAATCAAGATATGAGTGGAGACAYGGCATCCAGCARAGAAAGTTTGATATTTTRGATCAGGGGRMAAAAATAACCACTCGSACCATTMGSTATTCYTTGACATTTCGAACAGTTG | G | A | 930715 |

| | | | | |
|---|---|---|---|---|
| | TAAAAGATGCY | | | |
| chr5_15 | ACAGATAAAATCWTTCTAACATTTAAGTGAAAAAGTAATGGAGGTMC<br>AAGTTATAGAAGTAACRAAAGRAAATCTTGAGCYATTATGGAGTAAAA<br>TCGTGRAAGAYGYGAAAAACTCCATTTTYATTGCACTAGATACRGAA<br>ATGAGCGG[A/T]TTAGGACCCCAAGCMCAGCTWATGAAGTCTGACTT<br>GGASCAAAGGTATCARGGWATTTCTGCAGCAGCGTCWTCCCGTTCT<br>ATTATCTCTCTTGGACTTGCTTGTTTTGAGMCRTATTCTGGRAAAGAA<br>CCRCTGGCCTATAATGTAACT | A | T | 1055311 |
| chr5_16 | CACAAATATACCCAAATTGCCAGATTAGCGTCAAGTCGGCGTAATAG<br>CGACAGGAGGCTGCATGCTGGGTAYACAGACGAGAGTGCACCAGC<br>ATGGSAAGGGCATAGCGCGAAGAGAAGATTACGCAATAACTTAGAG<br>TACATGTGTGCG[C/T]CGCATATATTTCRTASTGYGGGGYCATGTGG<br>GATAYCTTTatcacctaaatcccatatttcttgatcgtgttttaaacacctgaAACCACTCCT<br>TTAGAGTGGTGGGGctacagttaaaaaattccttaaatattctttgtttactactt | C | T | 1078275 |
| chr5_18 | GCTCAAAATTAAGAGAAGGGATACTGCTTGAAGTACGGATTTGGGG<br>CGCCTAGTAAAGAGGGTCagaattaaacataatacattgtttttcTAGCTAAAAMC<br>GGACCAATAGTTTCAAAGGGTTAGGTTTAATATAGCATTACGAWATG<br>G[A/C]GCGTGTTCTTAAATTAWCGTTTTCTTTGAAATTTGTACAAAATA<br>GATTGAAATCCAATTGTGGTGTCGCGATATGACCAAAGCTTTAATATT<br>ATAACTGYCATGGCTTGTAATTTATGTCGCGGAACATAAAACCAGTAT<br>ATAATGATTA | A | C | 1221328 |
| chr5_19 | ARAAYGTAATTCAGCACATACTTMRAAACAGAGACAGCGTAATCATA<br>CATAACACATCAGTAGTTTGACCTTACATAACCYARCATATAACAAGA<br>CAGGAAGTGAATGTGGAAATTGCTATTTCTTCTTATTCAGCTGGACAT<br>RACACAG[T/C]TGCAGAAGGCCYATCTTAYCATAGCAATGCAATGTTA<br>TTGAATTggacaacaacaaaaWttgWaaagaTCAATGCYWCATAATTGccaR<br>WtttattaaagMaYATGATATYCACAATATTGCATGGTTGACTAACYTATTA<br>CTGCTAYGC | T | C | 1301011 |
| chr5_20 | gacgtaataagcaTGCsAwTGTGCAAGTCACAATGAATTTGTTGTCTGCAT<br>TGCGTCACAGTAAACCGATTGTTCGTGGAAAACkCGGTTAATGGTGT<br>CAACAACyAACAyGrAGCrAATGTATCGCGAAACTTGTCTAGTGGTTG<br>CGAk[T/C]GCGGAwGTTGGTCwACCTTGCTACTCGTGCGAAkGTAATT<br>ATCrGAATAATTyAGATTAACGCATCACGTTTAGTrTGTGTCGTCATAA<br>TGAArTCAyTGTCGGyTGCCAATCkkTATTTTrCATGTwACAACGAwTrrG<br>CAGTTCTGTC | T | C | 1400189 |
| chr5_21 | taaacaaacgAACCACGCGCCrAGATCTCACCACAGATAAGACGGTAAT<br>GGTTCGATGTGAGGTTGTCCAATCTCGCATTTTGCTTTGCATCGTCG<br>CGCTGCTATGTCAACCGCTwAGAAAACATTCAAwTTACAAAAGTTTCT<br>TTTTAC[A/T]ATTTCATAACGATCGACATAGCTTyATATTGCAGCAGTT<br>GTGTATTGCTAATGCAAGyGmATTAACCACACACAACTGCAsTGTTAT<br>CTAAAATGTTTACCATCCTTTCTTGGCAGTTTAAACTCCCCTGCGAAT<br>AAATAAGATATCTGT | A | T | 1475058 |

**Table S4: Cut-offs values based on VAFF values to determine homozygotes for each allele and heterozygotes as well as pure *C. intestinalis*, pure *C. robusta* and hybrids for the multiplex. Heterozygotes and hybrids had values in between the two cut-offs indicated in the table.**

| | **2012** | | **2021** | |
|---|---|---|---|---|
| **SNP** | **Cut-off homozygotes *C. intestinalis*** | **Cut-off homozygotes *C. robusta*** | **Cut-off homozygotes *C. intestinalis*** | **Cut-off homozygotes *C. robusta*** |
| mitochondrial | > 0.70 | < 0.40 | > 0.75 | < 0.30 |
| Chr5_SB | > 0.60 | < 0.40 | > 0.60 | < 0.30 |
| Chr5_10 | > 0.70 | < 0.40 | > 0.60 | < 0.30 |
| Chr5_11 | > 0.75 | < 0.4 | > 0.70 | < 0.30 |
| Chr5_12 | > 0.70 | < 0.40 | > 0.55 | < 0.30 |
| Chr5_13 | > 0.60 | < 0.40 | > 0.60 | < 0.40 |
| Chr5_15 | > 0.71 | < 0.40 | > 0.80 | < 0.30 |
| Chr5_16 | > 0.70 | < 0.35 | > 0.70 | < 0.35 |
| Chr5_18 | > 0.70 | < 0.40 | > 0.70 | < 0.40 |
| Chr5_19 | > 0.70 | < 0.40 | > 0.75 | < 0.40 |
| Chr5_20 | > 0.80 | < 0.50 | > 0.70 (run 1)<br>> 0.82 (run 2) | < 0.40 |
| Chr5_21 | > 0.60 | < 0.45 | > 0.60 | < 0.31 |
| | **Cut-off pure *C. intestinalis*** | **Cut-off pure *C. robusta*** | **Cut-off pure *C. intestinalis*** | **Cut-off pure *C. robusta*** |
| Multiplex (11 SNPs) | > 0.70 | < 0.40 | > 0.50 | < 0.30 |

**Table S5: p-values associated with A) Fisher's exact test on genotype counts per population and over the whole dataset at SNP 15 (core of the introgression) and B) Fisher's exact test and one-sided (2012 values greater than 2021) Wilcoxon signed-rank test on allele frequencies, for examining temporal changes in frequency of the foreign *C. robusta* allele, in the shoulders of the introgression island, between 2012 and 2021. p-values below 5% are indicated in bold.**

### A) Test for temporal changes at SNP 15 (i.e., the core of the introgression island)

| Site number | Site | p-value |
|---|---|---|
| 1 | Plymouth | 0.857 |
| 2 | Brixham | 0.857 |
| 3 | Southampton | 0.427 |
| 4 | Gosport | 0.856 |
| 6 | St-Vaast | 0.232 |
| 7 | St-Malo | NA |
| 8 | St-Quay | 1.00 |
| 9 | Perros Guirec | 0.340 |
| 10 | Roscoff | 0.679 |
| 11 | Aber Wrac'h | 0.126 |
| 12 | Brest | 0.367 |
| 13 | Camaret | 0.137 |
| 14 | Concarneau | 0.148 |
| 18 | La Rochelle | **0.049** |
| Over the whole dataset | | 0.494 |

### B) Test for temporal changes for each SNP located in the shoulders of the introgression island

| Locus | Fisher p-values | Wilcoxon p-values |
|---|---|---|
| chr5_10 | 0.655 | 0.639 |
| chr5_11 | 0.627 | 0.093 |
| chr5_12 | 0.058 | 0.099 |
| chr5_13 | **0.015** | **0.018** |
| chr5_16 | **0.005** | **0.002** |
| chr5_18 | **0.028** | **0.034** |
| chr5_19 | 0.680 | 0.091 |
| chr5_20 | **0.009** | **0.012** |
| chr5_21 | **$6\ 10^{-4}$** | **0.011** |
| Across all loci | **$3.41\ 10^{-6}$** | **$1.91\ 10^{-6}$** |

**Table S6: $r^2$ values for linkage disequilibrium between each SNP for both time periods, in 2012 (top) and 2021 (bottom).**

| 2012 | SNP10 | SNP11 | SNP12 | SNP13 | SNP15 | SNP16 | SNP18 | SNP19 | SNP20 |
|---|---|---|---|---|---|---|---|---|---|
| **SNP11** | 0.299 | | | | | | | | |
| **SNP12** | 0.122 | 0.087 | | | | | | | |
| **SNP13** | 0.060 | 0.037 | 0.564 | | | | | | |
| **SNP15** | 0.050 | 0.020 | 0.351 | 0.264 | | | | | |
| **SNP16** | 0.071 | 0.040 | 0.260 | 0.204 | 0.534 | | | | |
| **SNP18** | 0.064 | 0.039 | 0.191 | 0.177 | 0.313 | 0.432 | | | |
| **SNP19** | 0.002 | 0.000 | 0.019 | 0.025 | 0.093 | 0.104 | 0.375 | | |
| **SNP20** | 0.047 | 0.026 | 0.065 | 0.069 | 0.157 | 0.213 | 0.462 | 0.261 | |
| **SNP21** | 0.041 | 0.025 | 0.055 | 0.053 | 0.133 | 0.187 | 0.387 | 0.178 | 0.767 |

| 2021 | SNP10 | SNP11 | SNP12 | SNP13 | SNP15 | SNP16 | SNP18 | SNP19 | SNP20 |
|---|---|---|---|---|---|---|---|---|---|
| **SNP11** | 0.313 | | | | | | | | |
| **SNP12** | 0.197 | 0.102 | | | | | | | |
| **SNP13** | 0.113 | 0.047 | 0.498 | | | | | | |
| **SNP15** | 0.106 | 0.048 | 0.248 | 0.145 | | | | | |
| **SNP16** | 0.061 | 0.017 | 0.109 | 0.048 | 0.500 | | | | |
| **SNP18** | 0.028 | 0.007 | 0.078 | 0.047 | 0.149 | 0.270 | | | |
| **SNP19** | 0.000 | 0.000 | 0.012 | 0.002 | 0.060 | 0.132 | 0.377 | | |
| **SNP20** | 0.019 | 0.002 | 0.022 | 0.009 | 0.058 | 0.120 | 0.337 | 0.123 | |
| **SNP21** | 0.007 | 0.000 | 0.006 | 0.005 | 0.047 | 0.068 | 0.213 | 0.097 | 0.642 |
